# Cardiolipin deficiency disrupts electron transport chain and drives steatohepatitis

**DOI:** 10.1101/2024.10.10.617517

**Authors:** Marisa J. Brothwell, Guoshen Cao, J. Alan Maschek, Annelise M. Poss, Alek D. Peterlin, Liping Wang, Talia B. Baker, Justin L. Shahtout, Piyarat Siripoksup, Quentinn J. Pearce, Jordan M. Johnson, Fabian M. Finger, Alexandre Prola, Sarah A. Pellizzari, Gillian L. Hale, Allison M. Manuel, Shinya Watanabe, Edwin R. Miranda, Kajsa E. Affolter, Trevor S. Tippetts, Linda S. Nikolova, Ran Hee Choi, Stephen T. Decker, Mallikarjun Patil, J. Leon Catrow, William L. Holland, Sara M. Nowinski, Daniel S. Lark, Kelsey H. Fisher-Wellman, Patrice N. Mimche, Kimberley J. Evason, James E. Cox, Scott A. Summers, Zach Gerhart-Hines, Katsuhiko Funai

## Abstract

Metabolic dysfunction-associated steatotic liver disease (MASLD) is a progressive disorder marked by lipid accumulation, leading to metabolic dysfunction-associated steatohepatitis (MASH). A key feature of the transition to MASH involves oxidative stress resulting from defects in mitochondrial oxidative phosphorylation (OXPHOS). Here, we show that pathological alterations in the lipid composition of the inner mitochondrial membrane (IMM) directly instigate electron transfer inefficiency to promote oxidative stress. Specifically, mitochondrial cardiolipin (CL) was downregulated with MASLD/MASH in humans and in mice. Hepatocyte-specific CL synthase knockout (CLS-LKO) led to spontaneous and robust MASH with extensive steatotic and fibrotic phenotype. Loss of CL paradoxically increased mitochondrial respiratory capacity but also promoted electron leak primarily at sites III_QO_ and II_F_ of the electron transport chain, reduced the formation of I+III_2_+IV respiratory supercomplex, and disrupted the propensity of coenzyme Q (CoQ) to become reduced. Thus, low mitochondrial CL disrupts electron transport chain to promote oxidative stress and contributes to pathogenesis of MASH.

## Main

Metabolic dysfunction-associated steatotic liver disease (MASLD) is a growing global health concern with an increasing prevalence that parallels the rise in obesity.^1^ In the United States, annual medical costs related to MASLD exceed $103 billion.^2^ A large portion of patients with MASLD only exhibit steatosis, a silent and relatively benign early stage characterized by lipid accumulation in hepatocytes without hepatocellular inflammation.^3^ Steatosis can then progress to metabolic dysfunction-associated steatohepatitis (MASH), determined by hepatocyte injury and tissue fibrosis.^4^ MASH is the last stage of MASLD that may be reversible, making intervention at this stage particularly important.^3,5^ Although extensive clinical and basic research have been conducted in this field, the underlying mechanisms by which fatty liver transitions to MASH remain poorly understood.^6–8^

A defect in mitochondrial function is considered one of the hallmarks of MASLD progression in both mice and humans.^9–12^ MASLD is initially associated with an increase in mitochondrial respiratory capacity, followed by a subsequent impairment in oxidative phosphorylation (OXPHOS), and increased production of mitochondrial reactive oxygen species (ROS).^11,13^ Mitochondrial ROS is thought to be caused by an inefficient electron transport chain (ETC) that increases the propensity for electron leak. However, the mechanisms by which mitochondrial electron leak promotes MASLD are unknown.

Cardiolipin (CL) is a phospholipid with four acyl chains conjugated to two phosphatidylglycerol moieties linked by another glycerol molecule.^14^ CL resides almost exclusively in the inner mitochondrial membrane (IMM), constituting approximately 15-20% of the mitochondrial phospholipids.^15^ CL is synthesized by the condensation of phosphatidylglycerol (PG) and cytidine diphosphate-diacylglycerol (CDP-DAG) at the IMM via the enzyme cardiolipin synthase (CLS).^16,17^ Structural studies indicate that CL is essential for the activities of OXPHOS enzymes.^18–22^ In non-hepatocytes, decreased CL leads to compromised oxidative capacity,^23,24^ impaired membrane potential,^25^ and altered cristae morphology.^26^ In particular, low CL is associated with increased H_2_O_2_ production.^27,28^

In this manuscript, we set out to examine the changes in liver mitochondrial lipidome induced by MASH. In both mice and in humans, mitochondrial CL was downregulated in livers with MASLD/MASH compared to healthy controls. We then performed a targeted deletion of CLS in hepatocytes and studied its effects on liver, mitochondrial bioenergetics, and potential mechanisms that drive these changes.

## Results

### Mitochondrial cardiolipin levels are decreased in MASLD/MASH

Previous research from our lab in non-hepatocytes indicated that mitochondrial phospholipid composition affects OXPHOS electron transfer efficiency to alter electron leak.^15,29,30^ MASLD has been shown to alter the total cellular lipidome in liver.^31^ However, MASLD may also influence mitochondrial content in the hepatocytes, making it difficult to discern whether these are changes in the lipid composition of mitochondrial membranes and/or changes in cellular mitochondrial density. Thus, we performed liquid chromatography-tandem mass spectrometry (LC-MS/MS) lipidomics specifically on mitochondria isolated from human liver samples with or without MASH. Samples from patients undergoing liver transplant or resection due to end-stage MASH (Figure 1A, and Figure 1−table supplement 1) were compared to samples from patients without MASH, undergoing resection for benign liver tumors or metastases (Figure 1B, and Figure 1− figure supplement 1). Strikingly, mitochondrial CL levels were reduced, primarily due to a reduction in tetralinoleoyl-CL (72:8) (Figure 1C). This finding is consistent with previous studies demonstrating that MASLD is associated with reduced CL levels, and that tetralinoleoyl-CL is preferentially reduced in MASLD.^32–35^ Additionally, levels of mitochondrial PG, an essential precursor to CL production, were also reduced, suggesting a disruption in CL biosynthesis (Figure 1D-E).

**Figure 1.**
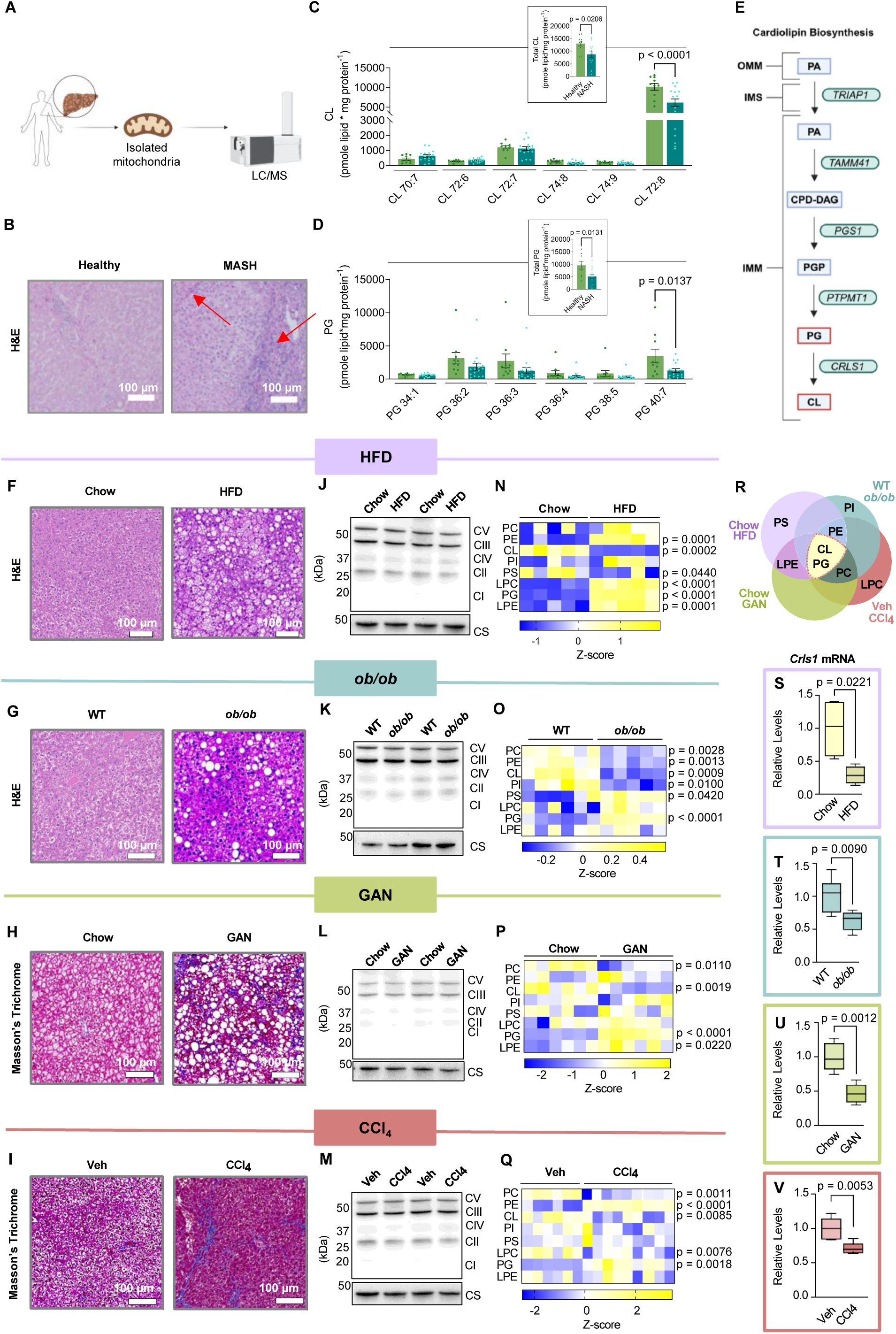
Hepatic mitochondrial phospholipidome in models of MASLD/MASH. (A) Schematic for mitochondrial phospholipidomic analyses in human liver samples. (B) Representative H&E staining of healthy or MASH livers. Arrows indicate fibrotic liver tissue. The diagnosis was based on obvious bridging fibrosis and/or regenerative nodules on H&E staining. (C, D) Hepatic mitochondrial CL and PG levels in healthy controls or individuals with MASH (*n*=10 and 16 per group). Data are shown as total FA chain length:number of double bonds. We have previously validated the 72:8 peak to be the tetralinoleoyl (18:2_18:2_18:2_18:2) cardiolipin on MS2, but we did not perform MS2 on every sample for the study. (E) Pathway for CL biosynthesis. (F–J) Representative H&E and Masson’s Trichrome staining of livers from mice under various MAFLD conditions: chow vs. Western HFD (16 wks), wildtype vs. ob/ob (20 wks), chow vs. GAN diet (30 wks), and vehicle vs. carbon tetrachloride (6 wks: 0.7 µL/g BW, twice weekly). (J–M) Western blot of OXPHOS subunits and citrate synthase in liver tissues from the same MAFLD models. (N–Q) Heatmaps of hepatic mitochondrial phospholipidomes from the same MAFLD models. (R) Venn diagram comparing hepatic mitochondrial phospholipidomes across all four MASLD models. (S–V) CLS mRNA levels in livers from the same MAFLD models (n=4–6 per group). Statistical significance was determined by 2-way ANOVA with within-row pairwise comparison (C, D, N–Q) and unpaired, two-sided Student’s t-test (C, D total lipid panels, S–V). All measurements were taken from distinct samples.

We next performed lipidomic analyses on four distinct mouse models of MASLD/MASH (Figure 1F-Q). These included: 1) mice given a Western high-fat diet (HFD, Envigo TD.88137) or standard chow diet for 16 weeks (Figure 1F), 2) ob/ob mice or their wildtype littermates at 20 weeks of age (Figure 1G), 3) mice given the Gubra Amylin NASH diet for 30 weeks (GAN, Research Diets D09100310) or standard chow (Figure 1H), 4) mice injected with carbon tetrachloride (CCI_4_) or vehicle (corn oil) for 6 weeks (Figure 1I). Importantly, none of these interventions appear to alter the protein abundances of OXPHOS subunits or citrate synthase (Figure 1J-M), suggesting that these interventions did not alter mitochondrial density in hepatocytes. Nevertheless, we performed all mitochondrial lipidomic analyses by quantifying lipids per mg of mitochondrial proteins. Each intervention appeared to alter different subsets of mitochondrial lipid classes (Figure 1N-Q, and Figure 1—figure supplement 2 and Figure 1—figure supplement 3), as seen with our previous studies in skeletal muscle and brown adipose tissues.^29,30^ We take these observations to mean that most physiological interventions induce multiple systemic and local responses that are not mechanistically directly related to the phenotype of interest (e.g., cold exposure or exercise can increase food intake, obesity could affect locomotion and insulation, etc.). Although several phospholipid classes were altered among the four models, strikingly, mitochondrial CL was reduced in all four MASLD/MASH models (Figure 1N-R). Furthermore, PG was significantly increased in all MASLD/MASH models (Figure 1N-R). These changes coincided with decreased transcript levels for CLS (Figure 1S-V). These observations further suggest that an insult in CL synthesis may be a key factor to disrupting mitochondrial function in MASLD/MASH.

### Hepatocyte-specific deletion of cardiolipin synthase promotes MASH

CL is thought to be exclusively synthesized in the IMM where CLS is localized. To study the role of CL in hepatocytes, we generated mice with hepatocyte-specific knockout of CLS (CLS-LKO for *CLS l*iver *k*nock*o*ut, driven by albumin-Cre) (Figure 2A,B), which successfully decreased mitochondrial CL levels (Figure 2C, and Figure 2—figure supplement 1). Additionally, CLS knockout resulted in a significant increase in mitochondrial phosphatidylglycerol (PG).^36^ As the substrate for CLS-mediated reaction, the increase in PG level was expected, Nonetheless, we cannot exclude the possibility that elevated PG levels could contribute to the observed phenotype. Consistent with our previous studies in non-hepatocytes, CLS deletion does not completely reduce CL levels to zero, suggesting that CL generated in other tissues may be imported. Our results showed that decreased levels of CL did not significantly impact body weight, food intake, or body composition (Figure 2D,E,F) but resulted in significantly less liver mass (Figure 2G).

**Figure 2.**
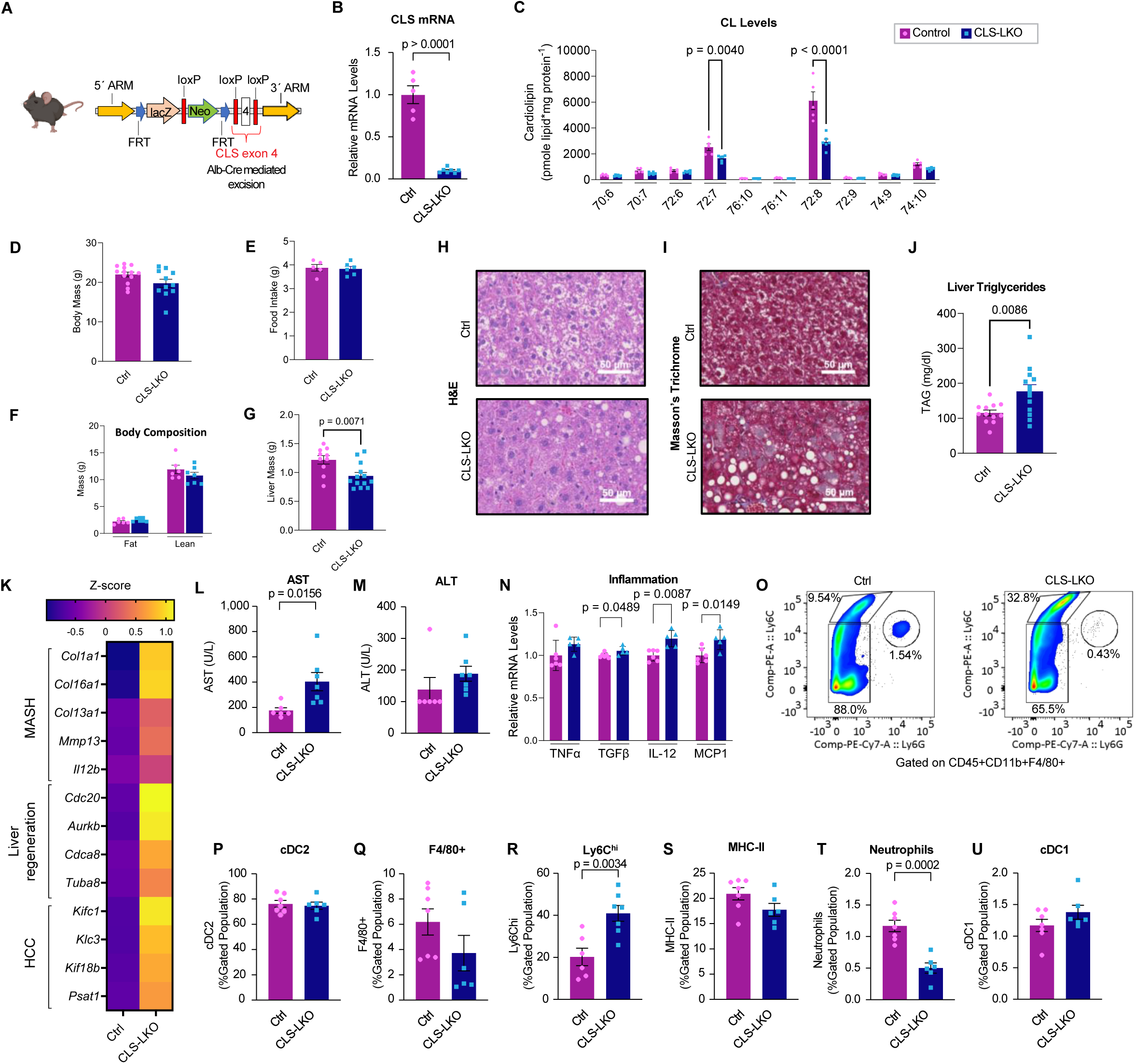
Hepatocyte-specific deletion of CLS induces MASLD/MASH. (A) Schematic for hepatocyte-specific deletion of CLS in mice. (B, C) CLS mRNA and mitochondrial CL levels in livers from control or CLS-LKO mice (n=5–7 per group). (D–G) Body mass, food intake, body composition, and liver mass of control or CLS-LKO mice (n=6–13 per group). (H, I) Representative H&E and Masson’s Trichrome staining of livers from control or CLS-LKO mice. (J) Liver triglycerides of control or CLS-LKO mice (n=12 per group). (K) RNAseq-derived heatmap of select genes associated with MASH, liver regeneration, and HCC (n=5–7 per group). (L, M) Serum AST and ALT levels (n=6–7 per group). (N) mRNA levels of TNFα, TGFβ, IL-12, and MCP1 in livers from control or CLS-LKO mice (n=5–7 per group). (O) Representative flow cytometry gating for liver cell populations in control or CLS-LKO mice. (P–U) Quantification of cDC2, F4/80+, Ly6C^hi^ inflammatory monocytes, MHC-II+, neutrophils, and cDC1 cell populations in livers (n=5–7 per group). All results are from mice fed standard chow. Statistical significance was determined by 2-way ANOVA with within-row pairwise comparison (C, N) and unpaired, two-sided Student’s t-test (B, D–G, K [adjusted for FDR], and P–U). Data represent mean ± SEM (B–G, L–N, P–U). All measurements were taken from distinct samples.

We sought to further characterize livers from control and CLS-LKO mice. Histological analyses revealed that CLS deletion was sufficient to promote steatosis, as evidenced by increased hepatic lipid accumulation and significantly elevated liver triglyceride levels (FigureL2H,LJ), as well as fibrosis (FigureL2I) under both standard chow-fed and high-fat-fed conditions (Figure 2—figure supplement 2A,B). To more comprehensively describe the effects of loss of hepatic CLS on gene expression, we performed RNA sequencing on these livers. CLS deletion increased the expression of 713 genes and decreased 1026 genes (Figure 2—figure supplement 2C). Pathway analyses revealed that many of the signature changes that occur with MASLD/MASH also occurred with CLS deletion (Figure 2K, and Figure 2—figure supplement 2D). This MASLD/MASH phenotype in our CLS knockout model was further confirmed with an elevation of the liver enzymes AST and ALT (Figure 2L,M) as well as increased mRNA levels of inflammatory markers (Figure 2N). We then proceeded to confirm these data by further phenotyping liver tissues from control and CLS-LKO mice.

In steatohepatitis, immune cell populations in the liver become altered to activate pathological immune response.^37^ Flow cytometry on livers from control and CLS-LKO mice indicated that the loss of CL promotes a robust classic immune response found in MASH (Figure 2O, and Figure 2—figure supplement 2E). cDC2 cells are a broad subset of dendritic cells with specific surface markers (e.g., CD11b, CD172a) that allow them to be distinguished from other dendritic cell populations.^38^ This broad population of dendritic cells was not different between control and CLS-LKO mice (Figure 2P). Notably, there was a marked reduction in the Kupffer cell population (Figure 2Q) - traditionally involved in maintaining liver homeostasis whose dysfunction can lead to dysregulated immune response.^39^ This reduction appears to be counterbalanced by a concomitant increase in Ly6C^hi^ population, which are known to typically go on to become inflammatory monocytes (Figure 2R). The replacement of Kupffer cells with other inflammatory cell populations suggests a shift towards a more pro-inflammatory environment, which may exacerbate liver injury and promote fibrosis. Nonetheless, the MHC-II cell population and neutrophils were not increased (Figure 2S,T) with neutrophils actually decreased (Figure 2T). The cDC1 cell population was not different (Figure 2U), which is traditionally elevated in response to cytotoxic T cells and might not be directly related to liver fibrosis.^40^ Together, these findings suggest that even on a chow diet, CLS-deficient livers exhibit inflammatory cell infiltration, a hallmark often associated with early signs of MASH.

### CLS deletion promotes fatty liver but increases mitochondrial respiratory capacity

Hepatocyte lipid accumulation may suggest defects in substrate handling, which is often manifested in systemic substrate handling. CLS deletion appeared to modestly improve glucose handling at select time points, but the magnitude of change was small and may reflect compensatory metabolic adaptations rather than a true improvement in systemic glucose homeostasis (FigureL3A-B). Effects on pyruvate handling were variable and did not show consistent directionality (FigureL3C-D). Lipid accumulation in hepatocytes can occur due to an increase in lipogenesis, a decrease in VLDL secretion, a decrease in β-oxidation, or increase in fatty acid uptake. However, mRNA levels for lipogenesis genes trended lower (not higher), and mostly unchanged for VLDL secretion, β-oxidation, or fatty acid uptake (Figure 3E, and Figure 3—figure supplement 1A). Circulating triglycerides were not lower in CLS-LKO mice compared to control mice (Figure 3—figure supplement 1B).

**Figure 3.**
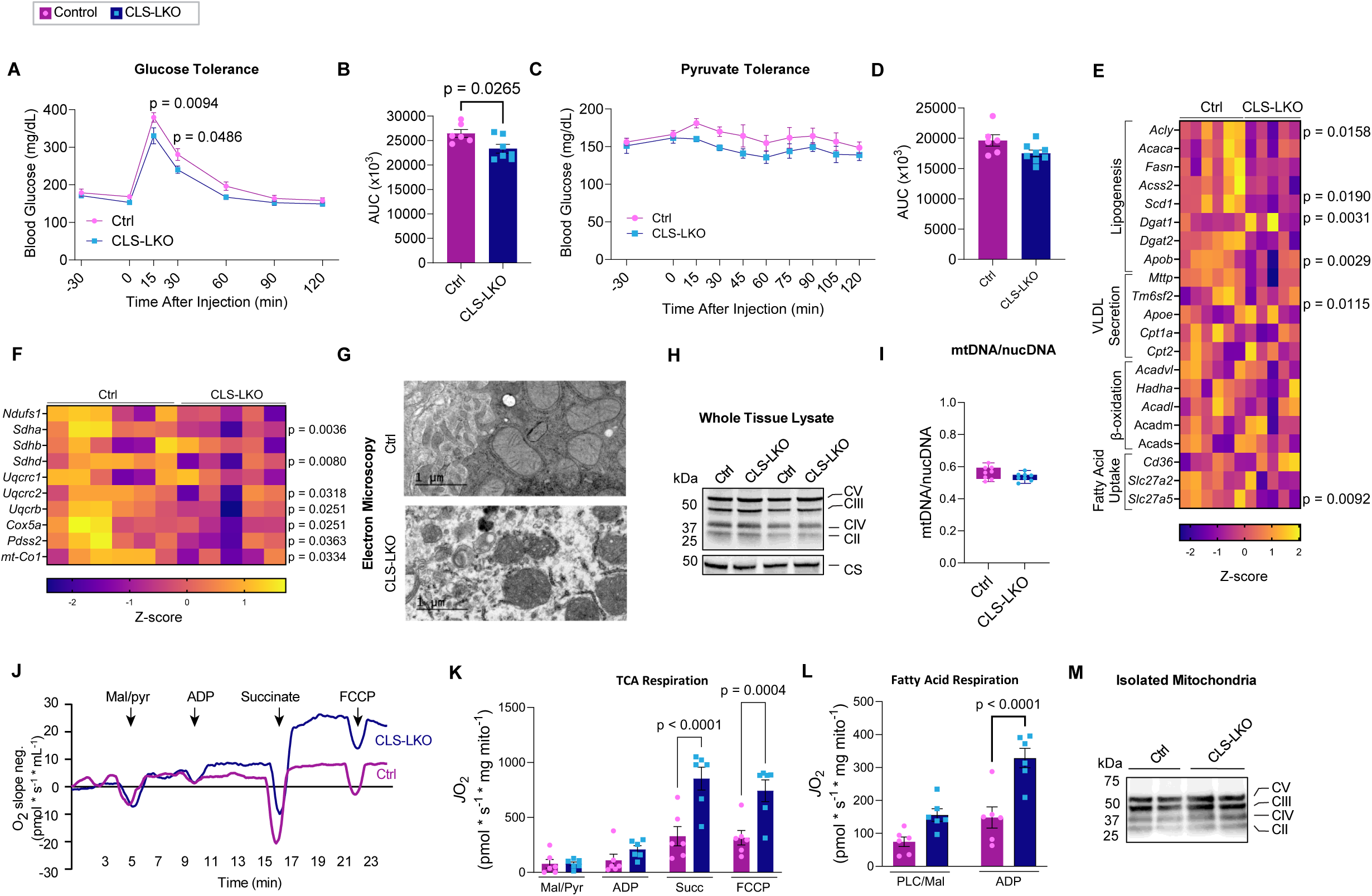
CLS deletion increases mitochondrial respiratory capacity. (A, B) Glucose tolerance test (IPGTT) and area under the curve (AUC) for control or CLS-LKO mice fed standard chow (n=6–7 per group). (C, D) Pyruvate tolerance test (PTT) and AUC (n=6–8 per group). (E, F) RNAseq-derived heatmaps of genes associated with lipogenesis, VLDL, β-oxidation, and ETS complex structure/function (n=5–6 per group). (G) Representative electron microscopy images of liver mitochondria from control or CLS-LKO mice. (H) Western blot of whole liver lysate using OXPHOS cocktail and citrate synthase for control or CLS-LKO mice. (I) Mitochondrial-to-nuclear DNA ratio in liver tissue from control or CLS-LKO mice (n=8 per group). (J) Representative tracing from high-resolution respirometry during maximal respiration using TCA cycle intermediates. (K, L) *J*O_2_ consumption in isolated liver mitochondria from control or CLS-LKO mice in response to malate, pyruvate, ADP, succinate, FCCP (K), or palmitoyl-carnitine, malate, and ADP (L) (n=6 per group). (M) Western blot of isolated mitochondria from livers of control or CLS-LKO mice using OXPHOS cocktail. All results are from mice fed standard chow. Statistical significance was determined by 2-way ANOVA with within-row pairwise comparison (A, C, K, L) and unpaired, two-sided Student’s t-test (B, D, E, F [adjusted for FDR], and I). Data represent mean ± SEM (A–D, K, L). Data in box-and-whiskers plot (I) represent median with min-to-max. All measurements were taken from distinct samples.

MASLD/MASH is known to be associated with reduced mitochondrial oxidative capacity, and such an effect may also occur with CL deficiency to induce lipid accumulation. Indeed, mRNA levels of several genes in the ETC were downregulated with CLS deletion, particularly those associated with structural components of the ETC complexes and the electron carrier CoQ (Figure 3F). Given that CL is located in the IMM where it binds to enzymes involved in OXPHOS,^41–44^ we reasoned that the loss of CL could reduce mitochondrial oxidative capacity to promote steatosis. Consistent with subcellular localization of CL, CLS deletion resulted in mitochondria with disorganized membrane structures and poorly developed cristae (Figure 3G). However, mitochondrial density quantified with western blots for respiratory complex subunits and citrate synthase (Figure 3H), as well as mtDNA/nucDNA (Figure 3I), showed no differences in livers from control and CLS-LKO mice. We thus speculated that CL lowers respiratory capacity not by reducing the total number of mitochondria or OXPHOS respirasomes, but by reducing the activity of respiratory enzymes. To our surprise, CLS deletion increased, rather than decreased, mitochondrial respiration (*J*O_2_), as measured by high-resolution Oroboros respirometry (Figure 3J), using both with Krebs cycle substrates (Figure 3K) as well as fatty acyl substrates (Figure 3L), to a similar degree. Importantly, these changes occurred in the absence of differences in OXPHOS subunit abundance per unit of mitochondria (Figure 3M), ruling out the possibility that changes in the abundance of respiratory enzymes are contributing to the change in respiration. A caveat to these findings is that CLS deletion promotes reduction in respiratory capacity after HFD-feeding (Figure 3—figure supplement 1C,D) However, CLS-LKO mice are steatotic in standard chow-fed condition, indicating that reduced mitochondrial fatty acid oxidation cannot be the cause of steatosis at baseline. The initial increase in respiration followed by its subsequent decrease is reminiscent of what is thought to occur with liver’s mitochondrial respiration over the course of MASLD progression.^45^

High-resolution respirometry experiments were performed in isolated mitochondria from hepatocytes by providing exogenous supraphysiological concentrations of substrates. While these assays provide robust measurements of respiratory capacity (the potential of mitochondria), they do not necessarily reflect their endogenous activity. To address this point, we performed stable isotope tracing experiments using uniformly labeled ^13^C-palmitate (Figure 4A)^46^ in murine hepa1-6 cells without (scrambled shRNA or shSC) or with CLS knockdown (shRNA targeting CLS or shCLS). Surprisingly, but in alignment the *J*O_2_ data, CLS deletion resulted in an increased fractional contribution of palmitate-derived carbon to TCA intermediates (Figure 4B-D), suggesting a shift towards enhanced fatty acid utilization. We also performed a similar tracing experiment using uniformly labeled ^13^C-glucose (Figure 4E-J, and Figure 4—figure supplement 1A-E). With CLS knockdown, we observed a shift in glucose-derived carbon incorporation, with a greater fraction of labeled carbon directed towards pyruvate (Figure 4E,F) and a smaller fraction towards lactate and alanine (Figure 4G, and Figure 4—figure supplement 1D). The fractional contribution of glucose-derived carbon to TCA intermediates was largely unchanged, except for a decrease in the fraction directed towards succinate (Figure 4H-J, and Figure 4—figure supplement 1B,C,E). While we do not attempt to resolve the specific routes of carbon entry into the TCA cycle, the observed labeling reflects overall glucose contribution to central metabolism. Despite the altered substrate incorporation, a decrease in TCA flux does not appear to account for the steatotic phenotype observed with CLS deletion. It’s important to note that hepa1-6 cells are hepatoma cells, and some aspects of metabolic response observed here may be different in hepatocytes.

**Figure 4.**
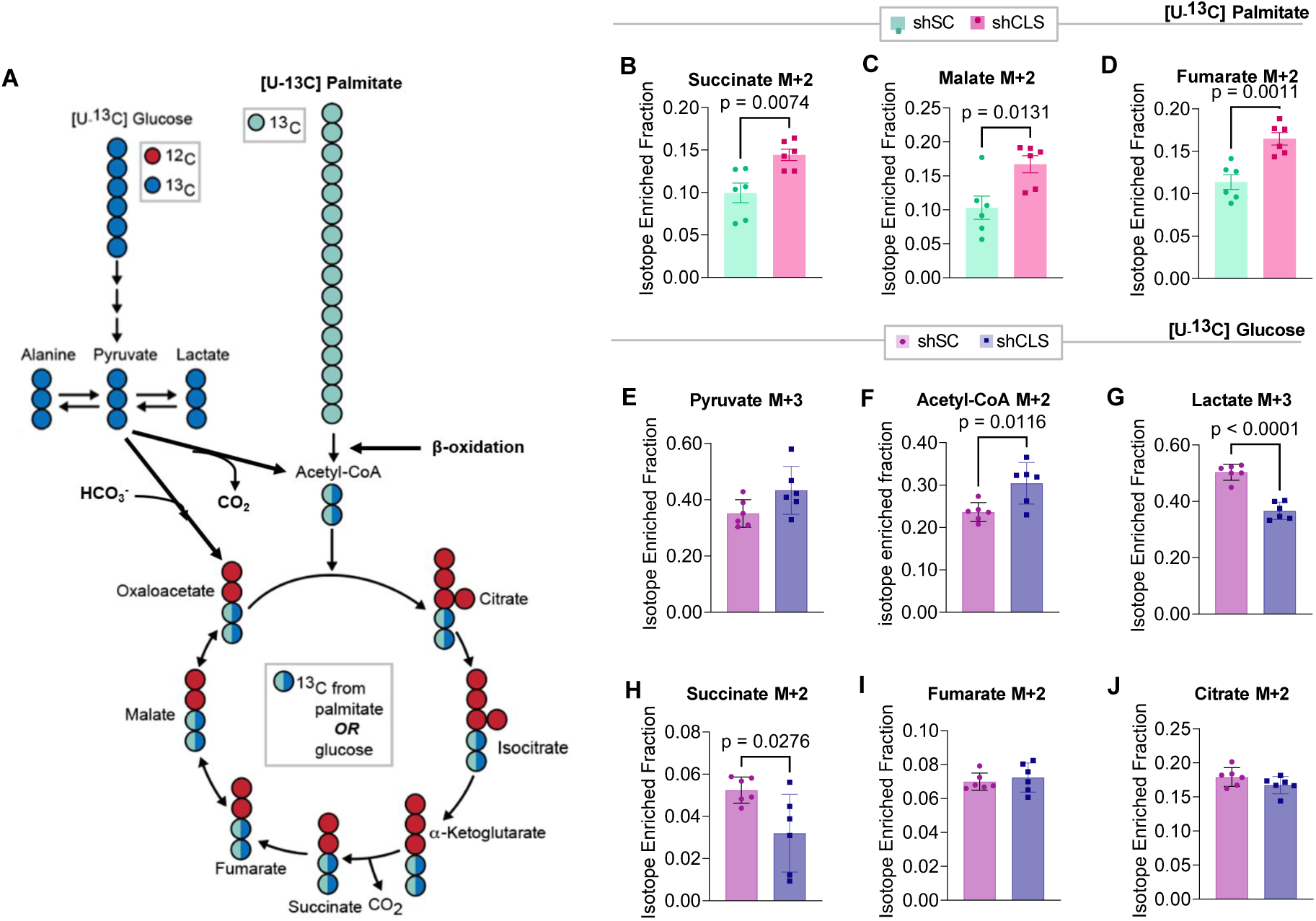
Stable isotope tracing with [U-^13^C] palmitate and [U-^13^C] glucose in hepa1-6 cells with or without CLS deletion. (A) Schematic of stable isotope tracing with [U-^13^C] palmitate or [U-^13^C] glucose, showing key intermediates in β-oxidation and the TCA cycle. (B–D) Levels of labeled succinate, malate, and fumarate from palmitate tracing in hepa1-6 cells without (shSC) or with CLS deletion (shCLS) (n=6 per group). (E–J) Levels of labeled pyruvate, acetyl-CoA, lactate, succinate, fumarate, and citrate from glucose tracing in hepa1-6 cells without or with CLS deletion (n=6 per group). Statistical significance was determined by unpaired, two-sided Student’s t-test. Data represent mean ± SEM. All measurements were taken from distinct samples.

### Low hepatic CL paradoxically increases mitochondrial respiratory capacity and promotes electron leak

Oxidative stress is thought to play a critical role in the transition from MASLD to MASH, wherein sustained metabolic insult leads to hepatocellular injury and collagen deposition resulting in fibrosis.^7^ CLS deletion promotes liver fibrosis in standard chow-fed condition (Figure 5A and Figure 2I) and in HFD-fed condition (Figure 2—figure supplement 2) that coincided with increased mRNA levels for fibrosis (Figure 5B and 2K). Tissue fibrosis is often triggered by apoptosis, and CLS deletion appeared to activate the caspase pathway (Figure 5C,D). How does deletion of CLS, a mitochondrial enzyme that produces lipids for IMM, activate apoptosis? Cytochrome c is an electron carrier that resides in mitochondria, which shuttles electrons between complexes III and IV.^47^ Under normal physiological conditions, cytochromeLc binds cardiolipin, which anchors it to the mitochondrial membrane.^44^ During the initiation of intrinsic apoptosis, CL can undergo oxidation and redistribution from the mitochondria to the cytosol. CL oxidation weakens its binding affinity for cytochrome c, releasing it from the IMM and into the OMM where it signals apoptosis.^13^ However, neither mitochondrial nor cytosolic cytochrome c abundance appeared to be influenced by CLS deletion (Figure 5E,F, and Figure 5—figure supplement 1A,B).

**Figure 5.**
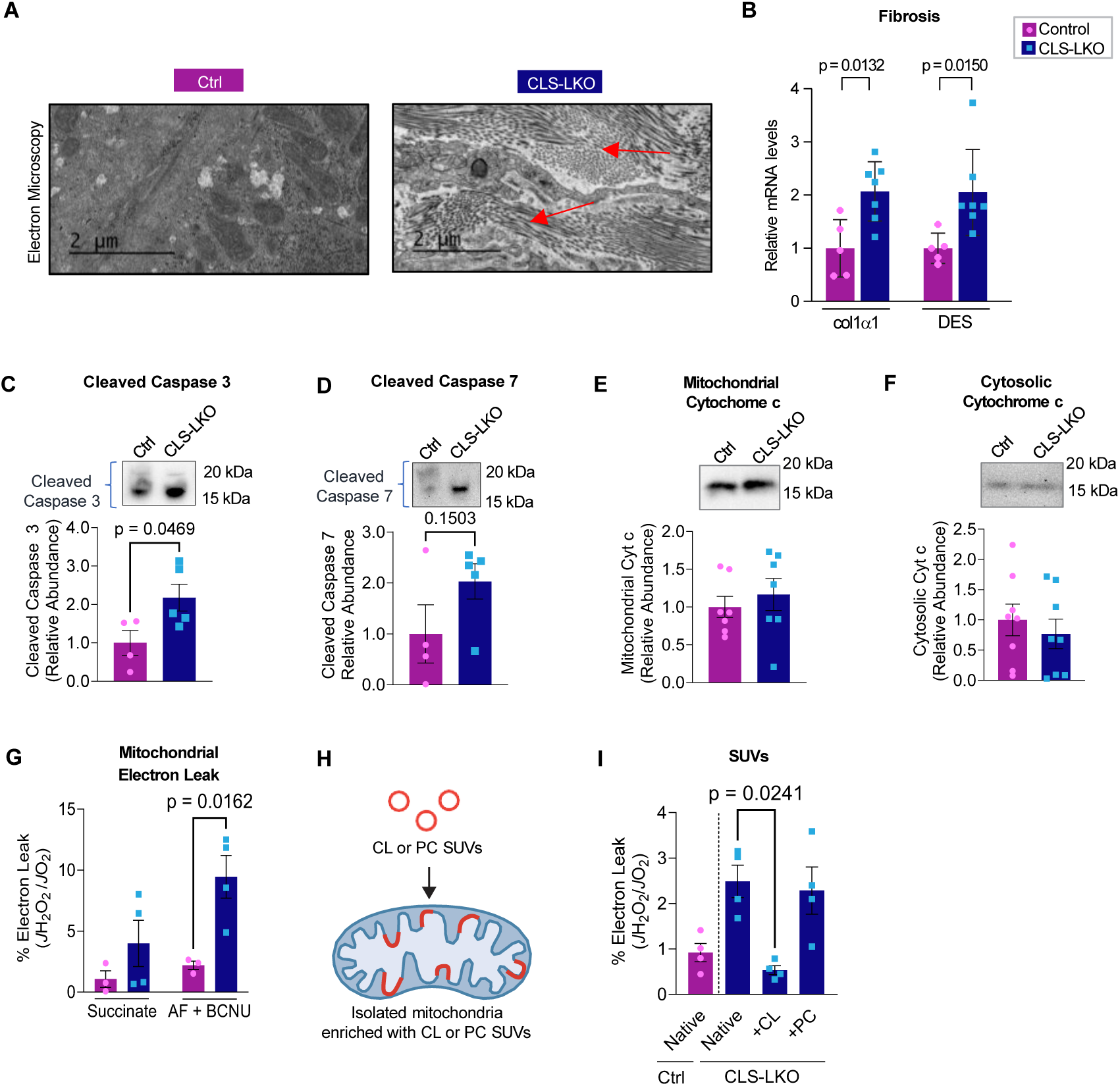
CL deficiency promotes mitochondrial electron leak. (A) Representative electron microscopy images of liver fibrosis in control or CLS-LKO mice (red arrows indicate collagen fiber bands arising from differing section orientations of the collagen/fibrotic matrix). (B) mRNA levels of fibrotic markers (Col1a1 and Desmin) in liver tissues (n=5–7 per group). (C–F) Levels of cleaved caspase-3, cleaved caspase-7, mitochondrial cytochrome c, and cytosolic cytochrome c in liver tissues (n=4–7 per group). (G) H_2_O_2_ emission in isolated liver mitochondria stimulated with succinate, or succinate with auranofin and BCNU (n=3–4 per group). (H) Schematic of small unilamellar vesicles (SUVs) containing cardiolipin (CL) or phosphatidylcholine (PC) for mitochondrial enrichment. (I) H_2_O_2_ production in liver mitochondria enriched with CL or PC SUVs (n=4 per group). All results are from mice fed standard chow. Statistical significance was determined by 2-way ANOVA with within-row pairwise comparison (B, G, I) and unpaired, two-sided Student’s t-test (C–F). Data represent mean ± SEM. All measurements were taken from distinct samples.

Mitochondrial ROS has been implicated in apoptosis and fibrosis with MASLD.^48–50^ Using high-resolution fluorometry in combination with high-resolution respirometry, we quantified electron leak from liver mitochondria with the assumption that almost all electrons that leak react with molecular O_2_ to produce O_2_^-^. Using recombinant superoxide dismutase, we ensure that all O_2_^-^ produced is converted into H_2_O_2_, which was quantified with the AmplexRed fluorophore.^51^ There were striking increases in mitochondrial electron leak in CLS-LKO mice compared to control mice on both standard chow (Figure 5G) and high-fat diet (Figure 5—figure supplement 1C). It is noteworthy that endogenous antioxidant pathways were insufficient to completely suppress oxidative stress induced by CLS deletion (H_2_O_2_ emission shown in the 1^st^ and 2^nd^ bars in Figure 5G, and Figure 5—figure supplement 1C). We also confirmed that *J*H_2_O_2_/*J*O_2_ was elevated with CLS knockdown in mitochondria from murine hepa1-6 cell line (Figure 5—figure supplement 1D) suggesting that low CL induces oxidative stress in a cell-autonomous manner.

While unknown, CLS may possess an enzymatic activity independent of CL synthesis that may contribute to electron leak. To more conclusively show that the loss of mitochondrial CL contributes to oxidative stress, we supplied exogenous CL to isolated mitochondria by fusing them with small unilamellar vesicles (SUVs) (Figure 5H).^52^ Isolated mitochondria from control and CLS-LKO mice were fused with SUVs containing either CL or phosphatidylcholine (PC) (Figure 5I, and Figure 5—figure supplement 1E). Remarkably, reintroducing CL to mitochondria from CLS-LKO mice reduced H_2_O_2_ production back to baseline, whereas PC had no effect. Thus, loss of CL drives the increased mitochondrial leak observed with CLS deletion.

### Loss of CL attenuates coenzyme Q interface with complex III

How does the loss of CL promote electron leak? We first addressed CL’s influence on the formations of respiratory supercomplexes. Respiratory supercomplexes exist in several combinations of multimers of Complex I, III, IV, and V and are thought to form either transiently or stably to improve electron transfer efficiency.^53,54^ CL may play an essential role in the stability of ETC supercomplexes.^55,56^ Using blue native polyacrylamide gel electrophoresis, we investigated supercomplex assembly in isolated hepatic mitochondria from control and CLS-LKO mice (Figure 6). When probing indiscriminately for all supercomplexes, there were many bands whose intensity were influenced by CLS deletion (Figure 6A). In particular, I+III_2_+IV supercomplex appeared to be substantially and consistently reduced in samples from CLS-LKO mice compared to control. To gain a more granular and definitive picture on the influence of CLS deletion on individual supercomplexes, we quantified supercomplexes using subunit-specific western blotting (Figure 6B-M). Strikingly, CLS deletion substantially reduced the abundance of the I+III₂+IV supercomplex when probing for UQCRFS1/complexLIII (FigureL6H,I), while increasing the levels of the CIII dimer (FigureL6I) and CIV monomer (FigureL6J,K), with a trend toward higher CI monomer abundance (FigureL6B-E). Although the functional implications of these alterations remain to be fully defined, disrupted supercomplex organization may underlie the elevated electron leak observed in CLS-deficient mitochondria. Together with our direct leak measurements, these data indicate that loss of CL compromises electron transport chain architecture, specifically by diminishing I+III₂+IV supercomplex assembly, which may promote electron leak.

**Figure 6.**
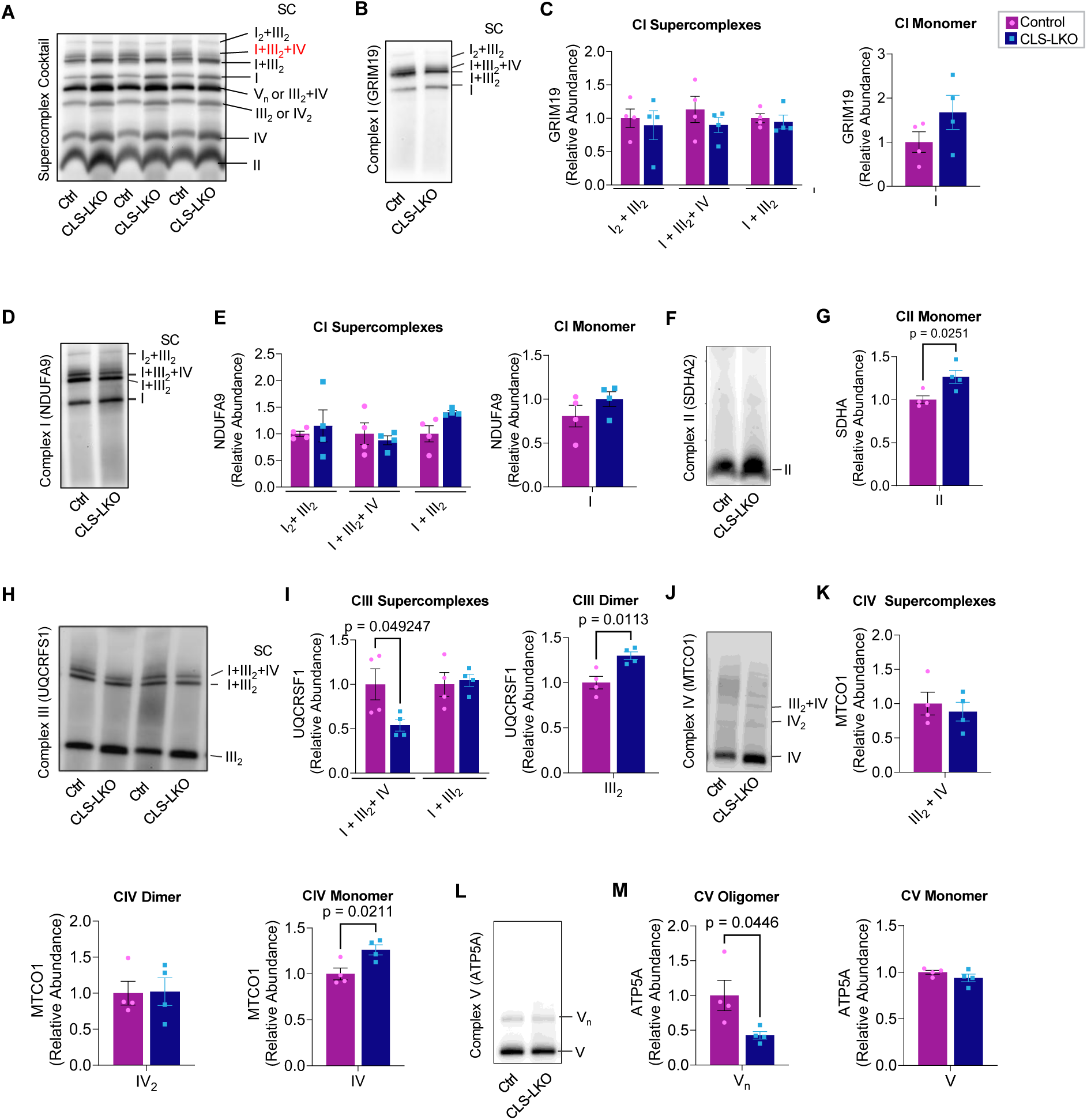
Influence of CL deficiency on respiratory supercomplex formation. (A–J) Representative western blots of respiratory supercomplexes in isolated liver mitochondria from control or CLS-LKO mice, detected using antibodies for supercomplexes, monomers, dimers, or oligomers, GRIM19/complex I (B), NDUFA9/complex I (D), SDHA2/complex II (F), UQCRFS1/complex III (H), MTCO1/complex IV (J), and ATP5A/complex V (L). Complex II is not thought to form respiratory supercomplexes. (C, E, G, I, K, M) Quantification of blots (n=4 per group). All results are from mice fed standard chow. Statistical significance was determined by 2-way ANOVA with within-row pairwise comparison (first panels in C, E, I, K) and unpaired, two-sided Student’s t-test (second panels in C, E, G, I, K, and both panels in M). Data represent mean ± SEM. All measurements were taken from distinct samples.

To better understand how low CL promotes inefficient electron transfer, we quantified electron leak at four specific sites of the ETC: 1) quinone-binding site in complex I (I_Q_), 2) flavin-containing site in complex I (I_F_), 3) succinate-dehydrogenase-associated site in complex II (II_F_), and 4) the ubiquinol oxidation site in complex (III_Qo_) Figure 7A). Electron leak at each of these sites can be quantified separately using substrates and inhibitors that restrict electron flow specific to these sites (Figure 7B-E). Among complex I, III, and IV (the components of I+III_2_+IV supercomplex), CLS deletion robustly increased electron leak at sites II_F_ and III_Qo_ (Figure 7D,E). In addition, electron leak either increased or trended to increase sites on complex I (Figure 7B,C). These findings raise a possibility that low CL may globally suppress the efficiency of the electron transport chain to promote electron leak.

**Figure 7.**
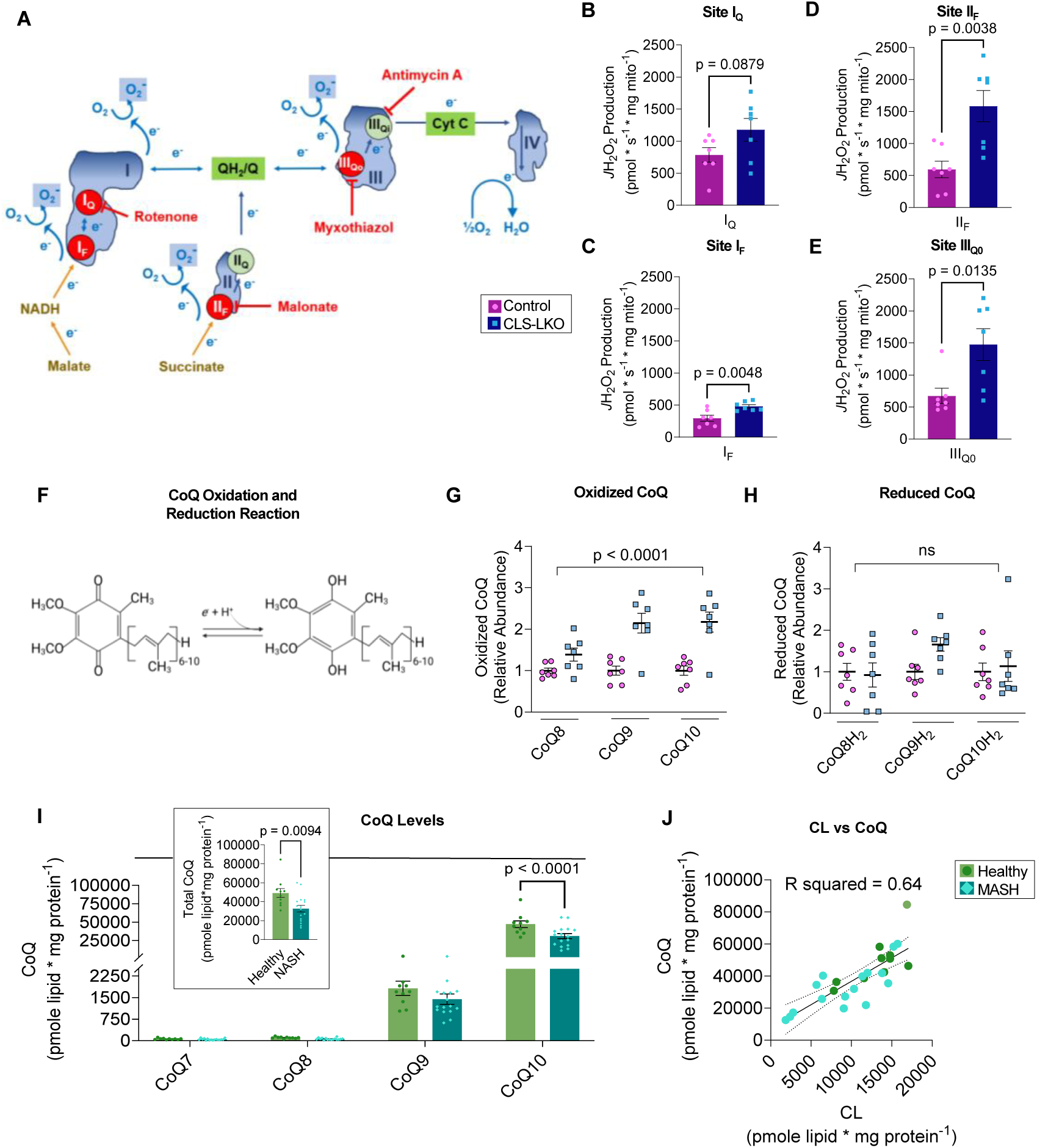
CL deficiency disrupts coenzyme Q homeostasis in humans and mice. (A) Schematic of site-specific electron leak. (B–E) Electron leak at sites I_Q_, I_F_, II_F_, and III_Q0_ in liver mitochondria from control or CLS-LKO mice (n=7 per group). (F) Oxidized CoQ (ubiquinone) can be reduced to CoQH_2_ (ubiquinol). (G, H) Oxidized and reduced CoQ levels in isolated liver mitochondria from control or CLS-LKO mice (n=7 per group). (I) CoQ levels in liver mitochondria from healthy human controls or patients with advanced steatohepatitis (n=10 and 16 per group). (J) Pearson correlation of CL and CoQ levels in human liver samples (R² = 0.64). Data in (B–E, G, H) are from mice fed standard chow. Statistical significance was determined by unpaired, two-sided Student’s t-test (B–E, panel of total levels in I) and 2-way ANOVA with within-row pairwise comparison (G, H, I). Data represent mean ± SEM (B–E, G, H, I). All measurements were taken from distinct samples.

To further examine the efficiency of the electron transport chain, we studied the redox state of CoQ, a carrier that feeds electrons from complex I and II into complex III. It is noteworthy that CoQ, like CL, is a lipid molecule (Figure 7F), and we thought it was conceivable that CL somehow interacts with CoQ to influence its electron transfer efficiency. Using redox mass spectrometry, we measured CoQ levels in whole liver tissues from control and CLS-LKO mice and found no difference in whole liver tissue CoQ levels (Figure 7—figure supplement 1A-I). However, since CoQ may also be found outside of mitochondria, we performed CoQ redox mass spectrometry in isolated mitochondria fractions of livers from control or CLS-LKO mice. Indeed, oxidized CoQ levels were increased (Figure 7G, and Figure 7—figure supplement 1J,L,N) in CLS-LKO mice compared to their controls. In contrast, reduced forms of CoQ were not lower in CLS-LKO mice compared to control mice (Figure 7H, and Figure 7—figure supplement 1K,M,O), and oxidized-to-reduced CoQ ratios in isolated mitochondria showed a similar pattern, with a significant increase observed only for total CoQ (Figure 7—figure supplement 1P,Q,R,S). Our findings that the normal level of reduced CoQ yielded greater oxidized CoQ, combined with increased electron flux (i.e., increase *J*O_2_, Figure 3J-L) can be interpreted to mean that there is a defect at the interface of CoQ-complex III that disrupts the efficient transfer of electrons promoting electron leak at site III_QO_. Conversely, it may also be interpreted that there are defects at complex I and II where oxidized CoQ are unable to become reduced due to lack of their ability to efficiently charge CoQ, also promoting electron leak.

To extrapolate these findings into humans, we went back to liver samples from patients undergoing liver transplant or resection due to end-stage MASH (Figure 1A) to quantify the abundance of mitochondrial CoQ. MASH was associated with a striking ∼40% reduction in the abundance of mitochondrial CoQ (Figure 7I) (tissue samples were not large enough to perform CoQ redox mass spectrometry on mitochondria). A Pearson correlation analysis showed a highly robust correlation between the abundances of mitochondrial CL and CoQ (Figure 7J, R^2^ = 0.64), indicating that the variability in the abundance of CL explains 64% of the variability of the abundance of CoQ. Based on our findings that CL is reduced with MASLD/MASH and that loss of CL influences CoQ electron transfer efficiency, we interpret these findings to mean that loss of CL destabilizes CoQ in humans.

## Discussion

In hepatocytes, disruptions of mitochondrial bioenergetics lead to and exacerbate metabolic-associated steatohepatitis.^57^ CL, a key phospholipid in the inner mitochondrial membrane, plays a critical role in mitochondrial energy metabolism.^23^ In this manuscript, we examined the role of CL in the pathogenesis of MASLD. In mice and in humans, MASLD/MASH coincided with a reduction in mitochondrial CL levels. Hepatocyte-specific deletion of CLS was sufficient to spontaneously induce MASH pathology, including steatosis and fibrosis, along with shift in immune cell populations towards a more pro-inflammatory profile, all of which occurred in mice given a standard chow diet. Paradoxically, high-resolution respirometry and stable isotope experiments showed that CLS deletion promotes, instead of attenuates, mitochondrial oxidative capacity in a fashion reminiscent of temporal changes that occur with mitochondrial bioenergetics in human MASH.^45^ Our principal finding on the role of hepatocyte CL in bioenergetics is that its loss robustly increases electron leak. This was likely due to: 1) direct interaction of CL with complex III, 2) lower abundance of I+III_2_+IV respiratory supercomplex, 3) global electron leak from the electron transport chain, and 4) loss of CL potentially interfering with the ability of CoQ to transfer electrons. In humans, mitochondrial CL and CoQ were co-downregulated in MASH patients compared to healthy controls, with a strong correlation (R² = 0.64) between CL and CoQ. Together, these results implicate CL as a key regulator of MASH progression, particularly through its effect on the electron transport chain to promote oxidative stress.

How might CL regulate CoQ? CoQ is the main electron transporter between complex I/II and III. CLS deletion in hepatocytes appeared to disrupt CoQ’s ability to cycle between its oxidized and reduced forms. There are several ways in which low CL might directly or indirectly influence CoQ’s redox state. The primary suspect is CL interacting with complex III, as eight or nine CL molecules are tightly bound to complex III^43^ and are thought to be essential to its function.^58^ While CL has been found to bind to other respiratory complexes, our data suggest that loss of CL might disproportionately influence complex III. This is also supported by our findings that the loss of CL reduced the formation of complex III-dependent supercomplex, without influencing other supercomplexes. Reduced capacity for complex III to efficiently accept electrons from CoQ might explain the increased electron leak at site III_Q0_ and increased level of oxidized CoQ. Meanwhile, loss of CL also increased electron leak at complex I and II. CL has also been implicated to bind to these complexes.^20,21,41^ Thus, complex III dysfunction is perhaps unlikely to entirely explain electron leaks at sites I_F_, I_Q_, and II_F_.^59^ Conversely, complex I and II are unlikely to be the only primary sites of defect as such defects probably will not promote electron leak at site III_QO_. Moreover, CLS deletion did not have an effect on the abundance of reduced CoQ despite there being greater abundance of oxidized CoQ, suggesting that complex I/II might also have reduced ability to transfer electrons onto CoQ. Another potential mechanism by which CL influences CoQ redox state is by CL directly interacting with CoQ. As they are both lipid molecules in the IMM, low CL may reduce the lateral diffusability of CoQ between respiratory complexes. Low CL might also indirectly influence CoQ by contributing to changes in membrane properties, distribution of ETC in the cristae, and the cristae architecture.^60^ Finally, increased electron leak, regardless of their origin, could have a feed-forward effect by which oxidative stress disrupts redox homeostasis in other components of ETC.

MASH is a progressive liver disease characterized by lipid accumulation, inflammation, and fibrosis in the liver.^61^ The progression to MASH involves a complex interplay of metabolic stress, mitochondrial defects, and immune responses that collectively promote hepatocellular injury.^62^ Our findings suggest that a low mitochondrial CL level directly induces key pathological features of MASH, including steatosis, fibrosis, and immune cell infiltration, even in the absence of dietary or environmental stressors, such as high-fat diet. When mice were fed a standard chow diet, CLS-LKO mice were more prone to lipid droplet accumulation than control mice. This phenotype was exacerbated when the mice were challenged with a high-fat diet. We primarily interrogated the mitochondrial bioenergetics of standard chow-fed control or CLS-LKO mice. A lower respiration rate would partially explain the lipid droplet accumulation, but to our surprise, CLS deletion increased *J*O_2_ regardless of substrates. Similarly, experiments using uniformly labeled ^13^C-palmitate or ^13^C-glucose showed that CLS deletion promoted an overall increase in the flux toward TCA intermediates, particularly for palmitate. CLS deletion did not appear to increase de novo lipogenesis or reduce VLDL secretion. Thus, it is unclear what mechanisms contribute to steatosis induced by CLS deletion.

Liver fibrosis is characterized by the accumulation of excess extracellular matrix components, including type I collagen, which disrupts liver microcirculation and leads to injury.^55^ Livers from CLS-LKO mice exhibited more fibrosis compared to control mice, even on a standard chow diet, which was worsened when fed a high-fat diet. Indeed, transcriptomic analyses revealed that CLS deletion activates pathways for fibrosis and degeneration, with many of the collagen isoforms upregulated. In the MASH liver, collagen deposition is accompanied by inflammatory cell infiltrate promoting an overall inflammatory environment. Flow cytometry experiments further confirmed that CLS deletion led to an increase in Ly6C^hi^ cell population, suggesting that dying resident Kupffer cells are being replaced by Ly6C^hi^ monocytes in the livers of CLS-LKO mice.^33^

Early in the MASLD disease progression, mitochondria adapt to the increased energy demands by increasing their respiratory capacity. In the later stages of disease progression to MASH, mitochondrial respiration diminishes.^63^ This pattern was reminiscent of our observations in the CLS-LKO mice. Livers from CLS-LKO mice fed a standard chow diet exhibited greater respiratory capacity compared to that of control mice. Conversely, livers from CLS-LKO mice on a high-fat diet exhibited lower respiratory capacity compared to that of control mice. We interpret these findings to suggest that liver mitochondria in chow-fed CLS-LKO mice are more representative of early stage of MASLD, while high-fat-diet fed CLS-LKO mice resemble later stages of MASLD.

In non-hepatocytes, low CL levels have been linked to electron leak in the context of a deficiency of the tafazzin gene, a CL transacylase, whose mutation promotes Barth syndrome.^15,26,64,65^ Paradoxically, we previously showed that CLS deletion in brown adipocytes does not increase electron leak.^30^ It is important to note that CLS deletion in our current or previous study does not completely eliminate CL (likely due to an extracellular source). We do not believe that CL is dispensable for efficient electron transfer in adipocytes. Rather, due to unclear mechanisms, different cell types likely exhibit varying tolerances to low CL influencing their bioenergetics, with brown adipocytes appearing more tolerant than hepatocytes. Regardless, electron leak was elevated with CLS deletion in both standard chow and high-fat diet-fed conditions. These observations mirror what has been shown in MASLD progression.^66^ The effect of CLS deletion on electron leak was due to reduced CL levels, as the reintroduction of cardiolipin via SUVs completely rescued the electron leak. While this rescue experiment provides strong evidence for the role of CL in regulating electron leak in hepatocytes, further studies are needed to determine whether CL reintroduction can rescue other aspects of the observed phenotype, such as steatosis, fibrosis, and altered glucose metabolism. Future experiments employing more comprehensive phenotyping following CL reintroduction will be valuable in addressing these questions.

CL is reported to be essential for the formation and stability of supercomplexes.^14,55,56,67^ CL has a distinctive conical shape with four fatty acyl chains, which allows it to create a highly curved membrane environment in the IMM, promoting close packing of protein complexes that likely facilitates supercomplex formation.^55^ CL also directly interacts with various subunits of the ETC complexes through electrostatic interactions, which help stabilize the supercomplexes by anchoring them together in a specific spatial orientation to optimize electron flow.^53^ Liver-specific deletion of CLS only resulted in a lower abundance of the I+III_2_+IV supercomplex. We interpret these findings to mean that I+III_2_+IV supercomplex is particularly sensitive to the reduced CL level in hepatocytes.

In our study, we observed a striking correlation between CL levels and CoQ in human liver samples from healthy/MASH patients (R_2_ of 0.64). In contrast, low mitochondrial CL induced by CLS deletion coincided with a greater mitochondrial CoQ content in CLS-LKO mice compared to controls. These data likely suggest that acute and robust reduction in CLS or CL level might trigger a compensatory CoQ production in mice. Conversely, because the samples from healthy/MASH patients were from those who had cirrhosis, MASH samples likely came from subjects that had suffered from years of MASLD pathology. In those samples, where reduction in CL was quantitatively modest compared to what was induced with CLS knockout, CoQ might have gradually decreased coincidental to the decrease in CL. Regardless of these differences in mice and humans, it is clear that there is a relationship between CL and CoQ that is worth further exploration.

In conclusion, our findings identify a critical role for CL in regulating ETC efficiency to promote oxidative stress. In both humans and mice, MASLD is associated with a decrease in hepatic mitochondrial CL, suggesting that low CL may be the cause of the obligatory increase in oxidative stress known to occur with MASLD progression. We further link CL deficiency to increased electron leak from the electron transport chain, and as the primary defect that promotes oxidative stress. We believe that these bioenergetic changes underlie the pathogenesis of MASLD, as CL deletion was sufficient to cause steatosis, fibrosis, and inflammation, phenocopying many changes that occur with MASLD/MASH progression. Further research will be needed to fully uncover how CL regulates CoQ, and to test whether rescuing the CL/CoQ axis might be effective in treating patients with MASLD/MASH.

## Methods

### Key resources table

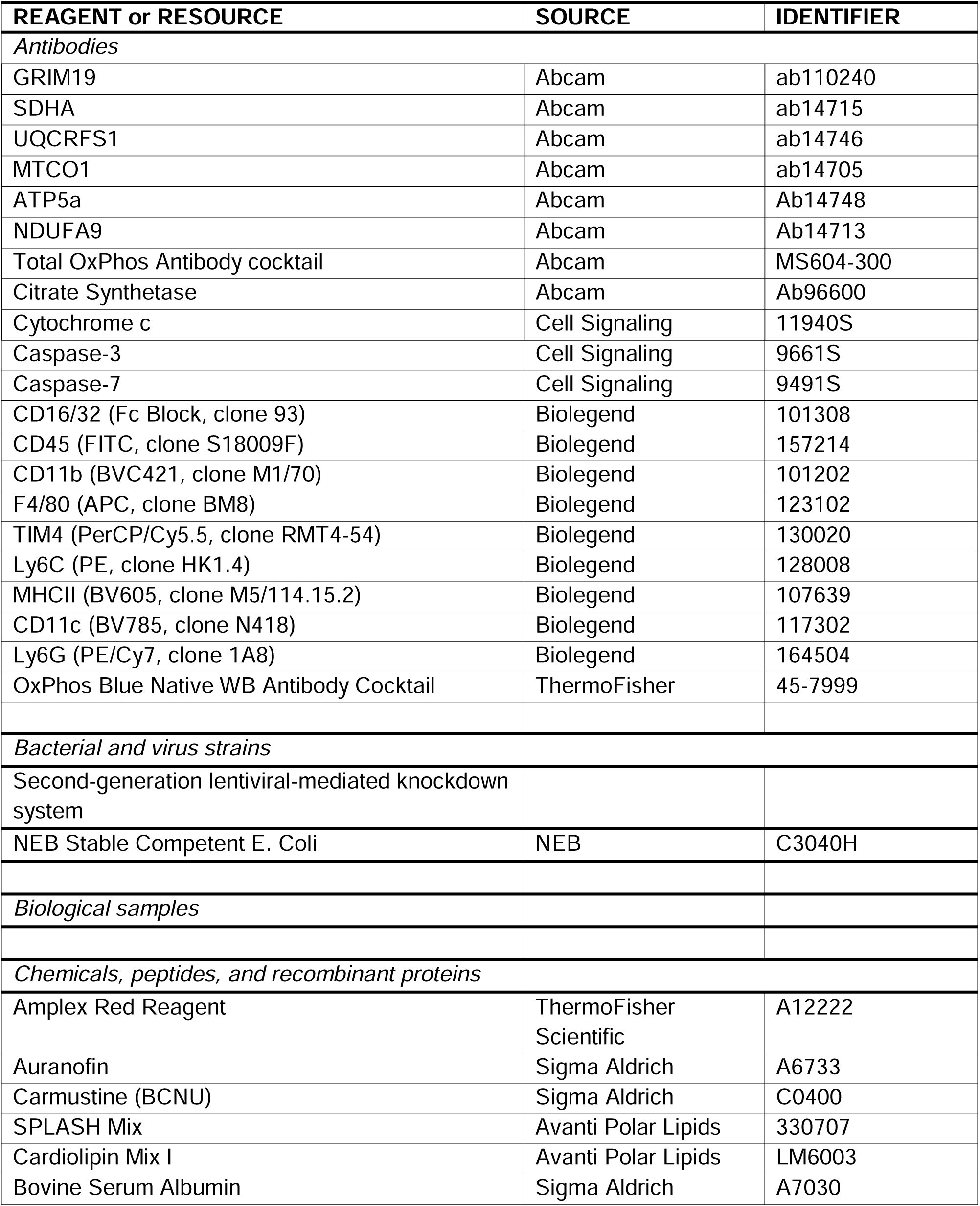
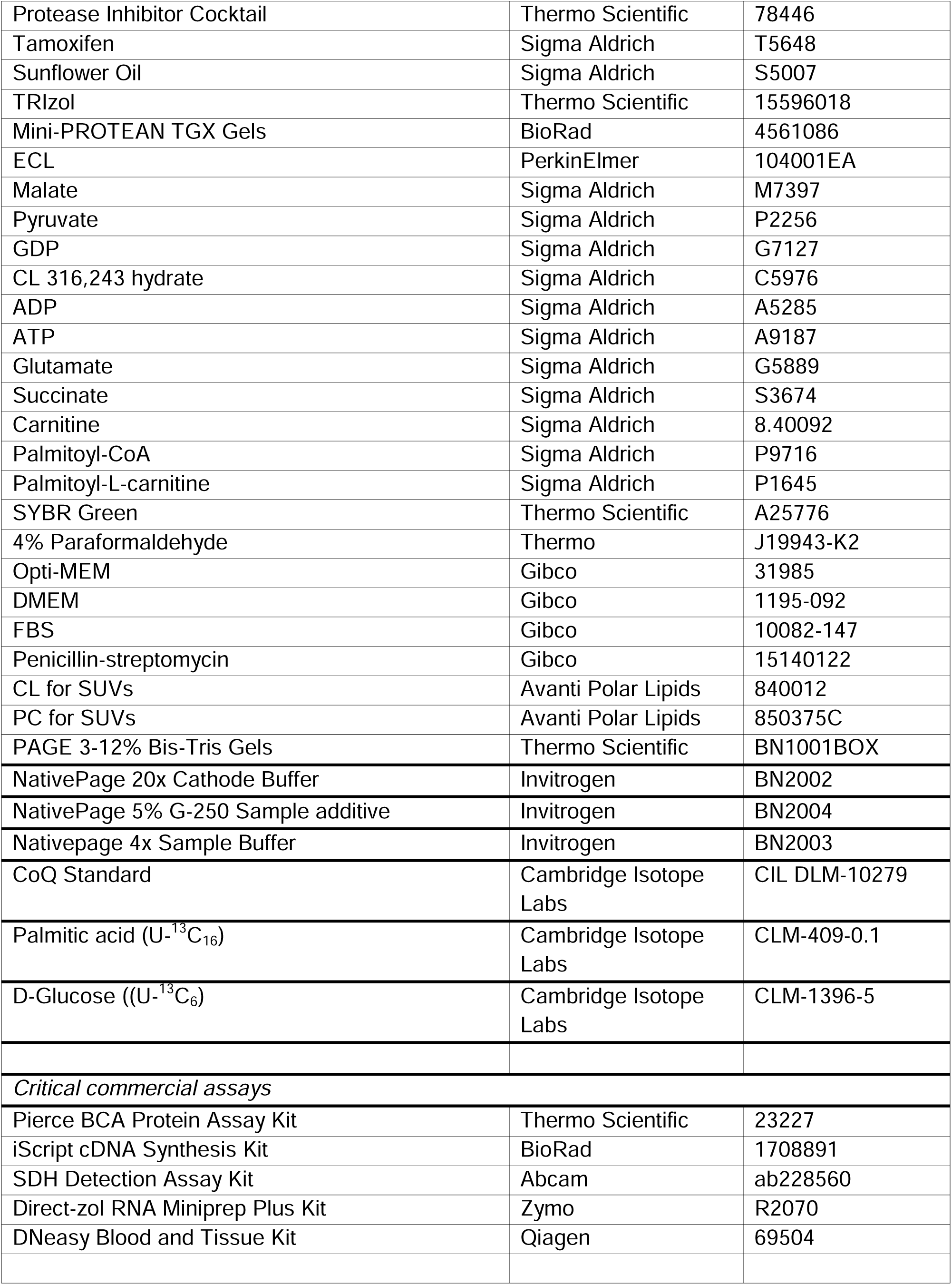
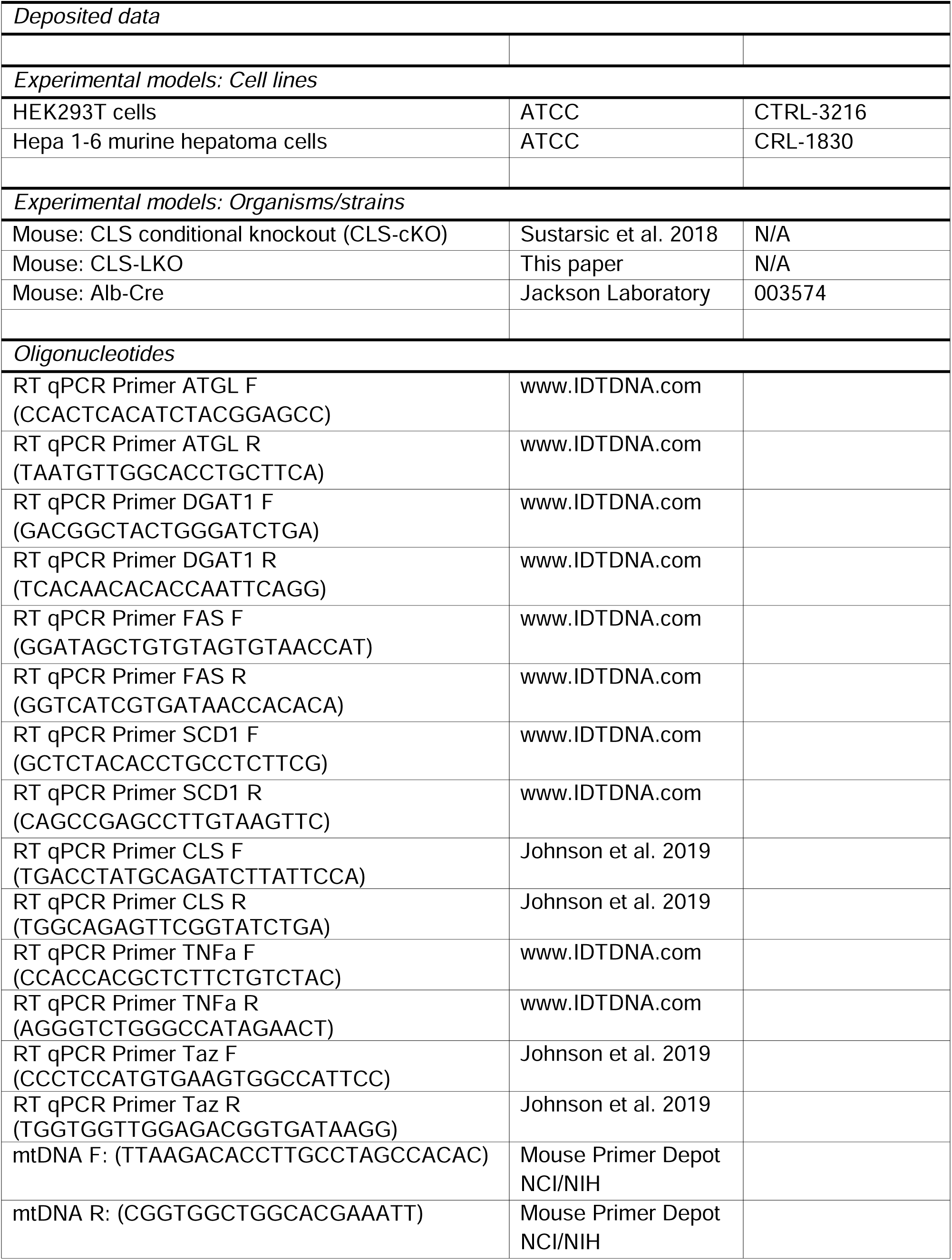
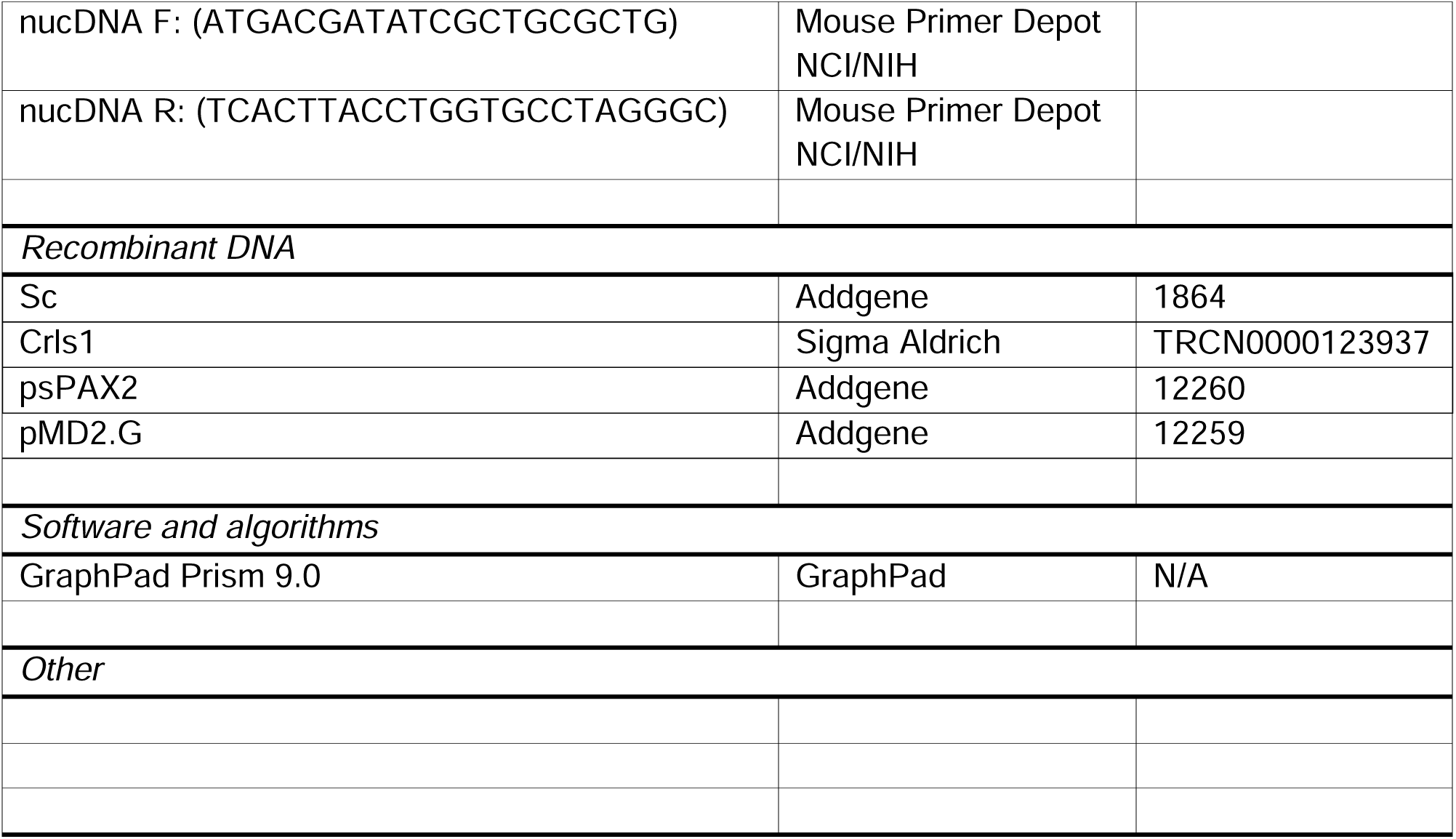

### Experimental model and subject details

#### Human participants

De-identified liver samples were acquired from the University of Utah Biorepository and Molecular Pathology Shared Resource from patients undergoing liver transplantation or resection due to end-stage liver disease and/or liver tumor(s). Informed consent was obtained from patients. All patients were classified to their respective diagnosis by a pathologist at the time of initial collection. The diagnosis for individual tissue samples was confirmed by a pathologist based on histology review of formalin-fixed, paraffin-embedded sections takes from the same location as the tissue analyzed by targeted lipid mass spectrometry.

#### Mice

All mice (male and female) used in this study were bred onto C57BL/6J background. CLS-LKO mice were generated by crossing the CLS conditional knockout (CLScKO^+/+^) generously donated by Dr. Zachary Gerhart-Hines (University of Copenhagen)^24^ with mice heterozygous for Albumin promotor Cre (Alb-Cre^+/-^) to produce liver-specific deletion of the CLS gene (CLScKO^+/+^, Alb-Cre^+/-^) or control (CLScKO^+/+^, no Cre) mice. CLScKO^+/+^ mice harbor loxP sites that flank exon 4 of the CLS gene. For high-fat diet studies, 8 wk CLS-LKO and their respective controls began high-fat diet (HFD, 42% fat, Envigo TD.88137) feeding for 8 wks. Mice were fasted 4 hours and given an intraperitoneal injection of 80 mg/kg ketamine and 10 mg/kg xylazine prior to terminal experiments and tissue collection. All animal experiments were performed with the approval of the Institutional Animal Care and Use Committees at the University of Utah.

#### Cell lines

Hepa 1-6 murine hepatoma cells were grown in high-glucose DMEM (4.5 g/L glucose, with L-glutamine; Gibco 11965-092) supplemented with 10% FBS (heat-inactivated, certified, US origin; Gibco 10082-147), and 0.1% penicillin-streptomycin (10,000 U/mL; Gibco 15140122). For lentivirus-mediated knockdown of CLS, CLS expression was decreased using the pLKO.1 lentiviral-RNAi system. Plasmids encoding shRNA for mouse *Crls1* (shCLS: TRCN0000123937) was obtained from MilliporeSigma. Packaging vector psPAX2 (ID 12260), envelope vector pMD2.G (ID 12259), and scrambled shRNA plasmid (SC: ID 1864) were obtained from Addgene. HEK293T cells in 10 cm dishes were transfected using 50 μL 0.1% polyethylenimine, 200 μL, 0.15 M sodium chloride, and 500 μL Opti-MEM (with HEPES, 2.4 g/L sodium bicarbonate, and l-glutamine; Gibco 31985) with 2.66 μg of psPAX2, 0.75 μg of pMD2.G, and 3 μg of either scrambled or *Crls1* shRNA plasmid. Cells were selected with puromycin throughout differentiation to ensure that only cells infected with shRNA vectors were viable.

### Method details

#### Body composition

To assess body composition, mice were analyzed using a Bruker Minispec NMR (Bruker, Germany) 1 week prior to terminal experiments. Body weights were measured and recorded immediately prior to terminal experiments.

#### RNA quantification

For quantitative polymerase chain reaction (qPCR) experiments, mouse tissues were homogenized in TRIzol reagent (Thermo Fisher Scientific) and RNA was isolated using standard techniques. The iScript cDNA Synthesis Kit was used to reverse transcribe total RNA, and qPCR was performed with SYBR Green reagents (Thermo Fisher Scientific). Pre-validated primer sequences were obtained from mouse primer depot (https://mouseprimerdepot.nci.nih.gov/). All mRNA levels were normalized to RPL32. For RNA sequencing, liver RNA was isolated with the Direct-zol RNA Miniprep Plus kit (Zymo Cat#: R2070). RNA library construction and sequencing were performed by the High-Throughput Genomics Core at the Huntsman Cancer Institute, University of Utah. RNA libraries were constructed using the NEBNext Ultra II Directional RNA Library Prep with rRNA Depletion Kit (human, mouse rat). Sequencing was performed using the NovaSeq S4 Reagent Kit v1.5 150×150 bp Sequencing with >25 million reads per sample using adapter read 1: AGATCGGAAGAGCACACGTCTGAACTCCAGTCA and adapter read 2: AGATCGGAAGAGCGTCGTGTAGGGAAAGAGTGT. Pathway analyses were performed by the Bioinformatics Core at the Huntsman Cancer Institute, University of Utah using the Reactome Pathway Database. For differentially expressed genes, only transcripts with Padj < 0.05 and baseMean > 100 are included.

#### DNA isolation and quantitative PCR

Genomic DNA for assessments of mitochondrial DNA (mtDNA) was isolated using a commercially available kit according to the manufacturer’s instructions (69504, Qiagen). Genomic DNA was added to a mixture of SYBR Green (Thermo Fisher Scientific) and primers. Sample mixtures were pipetted onto a 3840well plate and analyzed with QuantStudio 12K Flex (Life Technologies). The following primers were used: mtDNA fwd, TTAAGA-CAC-CTT-GCC-TAG-CCACAC; mtDNA rev, CGG-TGG-CTG-GCA-CGA-AAT-T; nucDNA fwd, ATGACG-ATA-TCG-CTG-CGC-TG; nucDNA rev, TCA-CTT-ACC-TGGTGCCTA-GGG-C.

#### Western blot analysis

For whole liver lysate, frozen liver was homogenized in a glass homogenization tube using a mechanical pestle grinder with homogenization buffer (50 mM Tris pH 7.6, 5 mM EDTA, 150 mM NaCl, 0.1% SDS, 0.1% sodium deoxycholate, 1% triton X-100, and protease inhibitor cocktail). After homogenization, samples were centrifuged for 15 min at 12,000 × g. Protein concentration of supernatant was then determined using a BCA protein Assay Kit (Thermo Scientific). Equal protein was mixed with Laemmeli sample buffer and boiled for 5 mins at 95°C for all antibodies except for OXPHOS cocktail antibody (at room temp for 5 mins), and loaded onto 4–15% gradient gels (Bio-Rad). Transfer of proteins occurred on a nitrocellulose membrane and then blocked for 1 hr. at room temperature with 5% bovine serum albumin in Tris-buffered saline with 0.1% Tween 20 (TBST). The membranes were then incubated with primary antibody (see Supplementary Files Table 2), washed in TBST, incubated in appropriate secondary antibodies, and washed in TBST. Membranes were imaged utilizing Western Lightning Plus-ECL (PerkinElmer) and a FluorChem E Imager (Protein Simple). For isolated mitochondria, identical procedures were taken with equal protein of mitochondrial preps.

#### Single cell preparation of liver tissue for flow cytometry

After mice were euthanized using isoflurane, blood was collected by cardiac puncture, the abdomen was exposed and the liver collected, rinsed with PBS and weighed. Liver was subsequently transferred in approximately 3ml of serum-free RPMI-1640 containing Collagenase D (10mg/ml; Sigma) and DNase (1mg/ml; Sigma) and incubated in a rocking platform for 45 min at 37°C. The liver extract was mashed through a 70µm filter and re-suspended in RPMI-1640 containing 10% FBS and centrifuged at 1600 rpm for 5 min. The supernatant was discarded and the pellet re-suspended in approximately 4 ml of 70% Percoll, then transferred to 15 ml conical tube, carefully overlayed with 4 ml of 30% Percoll and centrifuged 1600 rpm for 25 min with the brake turned off. The non-parenchymal cell suspension from the Percoll interface was removed and mixed with 10 mL of RPMI-1640 containing 10% FBS and the cells were centrifuged at 1600 rpm for 5 min. Red blood cells (RBC) were removed from the pelleted single cell suspensions of livers non-parenchymal cells by incubation in an ammonium chloride -based 1x RBC lysis buffer (Thermofisher, eBioscience). The cells were again pelleted and mixed with FACS buffer (2% BSA, 2mM EDTA in PBS), then stained with Zombie-NIR viability dye (BioLegend) per manufacturer’s instructions to discriminate live vs dead cells. To prevent non-specific Fc binding, the cells were incubated with Fc Block (anti-mouse CD16/32, clone 93, Biolegend) for 15 min followed by the indicated antibodies cocktail for 60 min in the dark on ice: CD45 (FITC, clone S18009F, Biolegend), CD11b (BVC421, clone M1/70, Biolegend), F4/80 (APC, clone BM8, Biolegend), TIM4 (PerCP/Cy5.5, clone RMT4-54, Biolegend), Ly6C (PE, clone HK1.4, Biolegend), MHCII (BV605, clone M5/114.15.2, Biolegend), CD11c (BV785, clone N418, Biolegend) and Ly6G (PE/Cy7, clone 1A8, Biolegend). After surface staining, cells were fixed with a paraformaldehyde-based fixation buffer (BioLegend). Flow cytometric acquisition was performed on a BD Fortessa X20 flow cytometer (BD Biosciences) and data analyzed using FlowJo software (Version 10.8.1; Tree Star Inc).

#### Glucose tolerance test

Intraperitoneal glucose tolerance tests were performed by injection of 1 mg glucose per gram body mass at least 6 days prior to sacrifice. Mice were fasted for 4 hours prior to glucose injection. Blood glucose was measured 30 minutes before glucose injection and at 0, 15, 30, 60, 90, and 120 minutes after injection via tail bleed with a handheld glucometer (Bayer Contour 7151H).

#### Pyruvate tolerance test

Pyruvate tolerance tests were performed by injection of 2 mg pyruvate per gram of body mass in PBS adjusted to pH 7.3-7.5 at least 6 days prior to sacrifice. Blood glucose was measured 30 minutes before pyruvate injection and at 0, 15, 30, 45, 60, 75, 90, 105, and 120 minutes after injection via tail bleed with a handheld glucometer (Bayer Contour 7151H).

#### Electron microscopy

To examine mitochondrial ultrastructure and microstructures, freshly dissected liver tissues from CLS-LKO and their controls were sectioned into ≈2 mm pieces and processed by the Electron Microscopy Core at University of Utah. To maintain the ultrastructure of the tissue via irreversible cross-link formation, each section was submerged in fixative solution (1% glutaraldehyde, 2.5% paraformaldehyde, 100 mM cacodylate buffer pH 7.4, 6 mM CaCl2, 4.8% sucrose) and stored at 4°C for 48 hours. Samples then underwent 3 × 10-minute washes in 100 mM cacodylate buffer (pH 7.4) prior to secondary fixation (2% osmium tetroxide) for 1 hour at room temperature. Osmium tetroxide as a secondary fixative has the advantage of preserving membrane lipids which are not preserved using aldehyde alone. After secondary fixation, samples were subsequently rinsed for 5 minutes in cacodylate buffer and distilled H_2_O, followed by prestaining with saturated uranyl acetate for 1 hour at room temperature. After prestaining, each sample was dehydrated with a graded ethanol series (2 × 15 minutes each: 30%, 50%, 70%, 95%; then 3 × 20 minutes each: 100%) and acetone (3 × 15 minutes) and were infiltrated with EPON epoxy resin (5 hours 30%, overnight 70%, 3 × 2-hour 40 minute 100%, 100% fresh for embed). Samples were then polymerized for 48 hours at 60°C. Ultracut was performed using Leica UC 6 ultratome with sections at 70 nm thickness and mounted on 200 mesh copper grids. The grids with the sections were stained for 20 minutes with saturated uranyl acetate and subsequently stained for 10 minutes with lead citrate. Sections were examined using a JEOL 1200EX transmission electron microscope with a Soft Imaging Systems MegaView III CCD camera.

#### Histochemistry

A fresh liver tissue was taken from each mouse and immediately submerged in 4% paraformaldehyde for 12 hours and then 70% ethanol for 48 hours. Tissues were sectioned at 10-μm thickness, embedded in paraffin, and stained for Masson’s Trichrome to assess fibrosis or hematoxylin and eosin (H&E) to determine fat droplet accumulation. Samples were imaged on Axio Scan Z.1 (Zeiss).

#### Native PAGE

Isolated mitochondria (100 µg) suspended in MIM were pelleted at 12,000 x g for 15 min and subsequently solubilized in 20 µL sample buffer (5 µL of 4x Native Page Sample Buffer, 8 µL 10% digitonin, 7 µL ddH_2_O per sample) for 20 min on ice and then centrifuged at 20,000 x g for 30 mins at 4°C. 15 µL of the supernatant (75 µg) was collected and placed into a new tube and mixed with 2 µL of G-250 sample buffer additive. Dark blue cathode buffer (50 mLs 20X Native Page running buffer, 50 mLs 20x cathode additive, 900 mLs ddH_2_O) was carefully added to the front of gel box (Invitrogen Mini Gel Tank A25977) and anode buffer (50 mLs 20x Native Page running buffer to 950 mL ddH_2_O) was carefully added to the back of the gel box making sure to not mix. The samples were then loaded onto a native PAGE 3-12% Bis-Tris Gel (BN1001BOX, Thermo Fisher Scientific), and electrophoresis was performed at 150 V for 1 hour on ice. The dark blue cathode buffer was carefully replaced with light blue cathode buffer (50 mLs 20X Native Page running buffer, 5 mL 20X cathode additive to 945 mLs ddH_2_O) and run at 30 V overnight at 4°C. Gels were subsequently transferred to PVDF at 100 V, fixed with 8% acetic acid for 5 min, washed with methanol, and blotted with the following primary antibodies Anti-GRIM19 (mouse monoclonal; ab110240), Anti-SDHA (mouse monoclonal; ab14715), Anti-UQCRFS1 (mouse monoclonal; ab14746), Anti-MTCO1 (mouse monoclonal; ab14705), Anti-ATP5a (mouse monoclonal; ab14748), Anti-NDUFA9 (mouse monoclonal; ab14713) in 5% non-fat milk in TBST. Secondary anti-mouse HRP antibody listed in the key resources table and Western Lightning Plus-ECL (PerkinElmer NEL105001) was used to visualize bands.

#### Isolation of mitochondrial and cytosolic fractions

Liver tissues were minced in ice-cold mitochondrial isolation medium (MIM) buffer [300 mM sucrose, 10 mM Hepes, 1 mM EGTA, and bovine serum albumin (BSA; 1 mg/ml) (pH 7.4)] and gently homogenized with a Teflon pestle. To remove excess fat in the samples, an initial high-speed spin was performed on all samples: homogenates were centrifuged at 12,000 x g for 10 mins at 4°C, fat emulsion layers were removed and discarded, and resulting pellets were resuspended in MIM + BSA. Samples were then centrifuged at 800 x g for 10 min at 4°C. The supernatants were then transferred to fresh tubes and centrifuged again at 1,300 x g for 10 min at 4°C. To achieve the mitochondrial fraction (pellet), the supernatants were again transferred to new tubes and centrifuged at 12,000 x g for 10 min at 4°C. The resulting crude mitochondrial pellets were washed three times with 0.15 M KCl to remove catalase, and then spun a final time in MIM. The final mitochondrial pellets were resuspended in MIM buffer for experimental use.

For cytosolic fraction, the supernatant from 12,000 x g spin was ultracentrifuged at 180,000 x g for 16 hours in a fixed angle rotor (FIBERLite, F50L-24×1.5). The top half of the supernatant was collected as analyzed as the cytosolic fraction for western blotting analyses. Mitochondrial protein content was determined by bicinchoninic acid assay using the Pierce BCA protein assay with bovine serum albumin as a standard.

#### Mitochondrial respiration measurements

Mitochondrial O_2_ utilization was measured using Oroboros O2K Oxygraphs. Isolated mitochondria (50 µg for TCA substrate respiration and 100 µg for fatty acid respiration) were added to the oxygraph chambers containing assay buffer Z (MES potassium salt 105 mM, KCl 30 mM, KH_2_PO_4_ 10 mM, MgCl_2_ 5 mM, BSA 1 mg/ml). Respiration was measured in response to the following substrates: 0.5 mM malate, 5 mM pyruvate, 2.5 mM ADP, 10 mM succinate, 1.5 μM FCCP, 0.02 mM palmitoyl-carnitine, 0.5 mM malate.

#### Mitochondrial *J*H_2_O_2_

Mitochondrial H_2_O_2_ production was determined in isolated mitochondria from liver tissue using the Horiba Fluoromax-4/The Amplex UltraRed (10 μM)/horseradish peroxidase (3 U/ml) detection system (excitation/emission, 565:600, HORIBA Jobin Yvon Fluorolog) at 37°C. Mitochondrial protein was placed into a glass cuvette with buffer Z supplemented with 10 mM Amplex UltraRed (Invitrogen), 20 U/mL CuZn SOD). Since liver tissue is capable of producing resorufin from amplex red (AR), without the involvement of horseradish peroxidase (HRP) or H_2_O_2_, phenylmethylsulfonyl fluoride (PMSF) was included to the experimental medium due to its ability to inhibit HRP-independent conversion of AR to resorufin.^68^ PMSF was added to the cuvette immediately prior to measurements and at a concentration that does not interfere with biological measurements (100 µM).^68^ A 5-min background rate was obtained before adding 10 mM succinate to the cuvette to induce H_2_O_2_ production. After 4 min, 100 µM 1,3-bis(2-chloroethyl)-1-nitrosourea (BCNU) was added to the cuvette with 1 µM auranofin to inhibit glutathione reductase and thioredoxin reductase, respectively. After an additional 4 min, the assay was stopped, and the appearance of the fluorescent product was measured.

Site-specific electron leak was measured by systematically stimulating each site while inhibiting the other three.^59^ Site I_F_ was investigated in the presence of 4 mM malate, 2.5 mM ATP, 5 mM aspartate, and 4 µM rotenone; site I_Q_ was measured as a 4 µM rotenone-sensitive rate in the presence of 5 mM succinate; site III_QO_ was measured as a 2 µM myxothiazol-sensitive rate in the presence of 5 mM succinate, 5 mM malonate, 4 µM rotenone, and 2 µM antimycin A; and site II_F_ was measured as the 1 mM malonate-sensitive rate in the presence of 0.2 mM succinate and 2 µM myxothiazol. As previously mentioned, electron leak is quantified using Amplex Red in the presence of excess superoxide dismutase, such that both superoxide and hydrogen peroxide production are accounted for by a change in fluorescence intensity (JH_2_O_2_) using high-resolution fluorometry (Horiba Fluoromax4®).

#### Lipid mass spectrometry for reduced and oxidized CoQ

Liver tissue was homogenized in ice cold isolation buffer (250 mM sucrose, 5 mM Tris-HCl, 1 mM EGTA, 0.1% fatty acid free BSA, pH 7.4, 4°C) using a tissuelyser. Mitochondria were then isolated via differential centrifugation (800 x g for 10 min, 1300 x g for 10 min, 12,000 x g for 10 min at 4°C), flushing each step under a stream nitrogen to prevent oxidation. Protein content was determined by bicinchoninic acid assay using the Pierce BCA protein assay with bovine serum albumin as a standard. To extract CoQ from mitochondria, incubations of 100 µg mitochondrial protein in 250 µL ice-cold acidified methanol, 250 µL hexane, and 1146 pmol per sample of CoQ standard (Cambridge Isotope Laboratories, CIL DLM-10279) were vortexed. The CoQ-containing hexane layer was separated by centrifugation (10 min, 17,000 x g, 4°C) and then dried down under a stream of nitrogen. Dried samples were then resuspended in methanol containing 2 mM ammonium formate and transferred to 1.5 mL glass mass spectrometry vials. Liquid chromatography-mass spectrometry (LC-MS/MS) was then performed on the reconstituted lipids using an Agilent 6530 UPLC-QTOF mass spectrometer.

#### U-^13^C-glucose and U-^13^C-palmitate labeling in hepa1-6 cell line

Stable isotope tracing experiments were conducted using U-^13^C-glucose or U-^13^C-palmitate to assess metabolic fluxes in hepa1-6 cells with or without CLS knockdown. Cells were incubated in either U-^13^C-glucose or U-^13^C-palmitate for 4 hours. For palmitate incubations, U-^13^C-palmitate was conjugated to fatty acid-free bovine serum albumin (BSA) and supplemented with 1 mM carnitine prior to being added to the culture medium. Following the 4-hour incubation period, cells were washed once with 1 mL of ice-cold Dulbecco’s Phosphate Buffered Saline (dPBS) and then scraped in 0.7 mL ice-cold dPBS. The cells were pelleted by centrifugation at 500 x g for 5 minutes at 4°C, and the supernatant was carefully removed. Cell pellets were resuspended in 215 µL of ice-cold dPBS and sonicated on ice using a 1/8” probe sonicator at 30% amplitude for 6 seconds. Debris was lightly pelleted by centrifugation at 500 x g for 2 minutes at 4°C. From the resulting supernatant, 25 µL was transferred to a separate tube for protein quantification using the bicinchoninic acid (BCA) assay. Samples incubated with U-^13^C-glucose were submitted for gas chromatography-mass spectrometry (GC-MS) and liquid chromatography-mass spectrometry (LC-MS) analysis, while those incubated with U-^13^C-palmitate were submitted for GC-MS analysis only.

#### GC-MS

All GC-MS analysis was performed with an Agilent 5977b GC-MS MSD-HES and an Agilent 7693A automatic liquid sampler. Dried samples were suspended in 40 µL of a 40 mg/mL O-methoxylamine hydrochloride (MOX) (MP Bio #155405) in dry pyridine (EMD Millipore #PX2012-7) and incubated for one hour at 37 °C in a sand bath. 25 µL of this solution was added to auto sampler vials. 60 µL of N-methyl-N-trimethylsilyltrifluoracetamide (MSTFA with 1% TMCS, Thermo #TS48913) was added automatically via the auto sampler and incubated for 30 minutes at 37 °C. After incubation, samples were vortexed and 1 µL of the prepared sample was injected into the gas chromatograph inlet in the split mode with the inlet temperature held at 250 °C. A 10:1 split ratio was used for analysis for most metabolites. Any metabolites that saturated the instrument at the 10:1 split was analyzed at a 100:1 split ratio. The gas chromatograph had an initial temperature of 60 °C for one minute followed by a 10 °C/min ramp to 325 °C and a hold time of 10 minutes. A 30-meter Agilent Zorbax DB-5MS with 10 m Duraguard capillary column was employed for chromatographic separation. Helium was used as the carrier gas at a rate of 1 mL/min.

Data were collected using MassHunter software (Agilent). Metabolites were identified and their peak area was recorded using MassHunter Quant. This data was transferred to an Excel spread sheet (Microsoft, Redmond WA). Metabolite identity was established using a combination of an in-house metabolite library developed using pure purchased standards, the NIST library and the Fiehn library. There are a few reasons a specific metabolite may not be observable through GC-MS.

#### LC-MS

A pooled quality control (QC) sample for each sample type was prepared by combining 10 µL of each sample. Samples were randomized prior to analysis. A SCIEX 7600 Zeno-ToF (AB SCIEX LLC, Framingham, MA, USA) coupled to an Agilent 1290 Infinity II HPLC system in positive-ionization modes was used for analysis. Separation was achieved using a Waters BEH zhilic 2.1 x 100 mm column (Waters Corporation, Milford, MA, USA) with Phenomenex Krudkatcher Ultra (Phenomenex, Torrence, CA, USA). Buffers consisted of 99% ACN with 5% ddH2O (buffer B) and 5% 25 mM ammonium carbonate in ddH2O (buffer A). An initial concentration of 99% buffer B was decreased to 85% over 2 min, then further decreased to 75%, and 60% over 3-minute intervals. Next, buffer B was decreased to 40% and held for 1 min. Then Buffer B was decreased to 1% and held for 1 min. Eluents were returned to initial conditions over 0.1 min, and the system was allowed to equilibrate for 5 min between runs. Mass spectrometry analysis was performed by high-resolution multiple reaction monitoring (MRM HR).

Data was collected using SCIEX analyst. Chromatogram integration was performed using SCIEX MultiQuant. This data was transferred to an Excel spread sheet (Microsoft, Redmond WA). Statistical analysis was performed using Microsoft Excel Data Analysis add-in. The metabolite lists were curated to the metabolites of interest for the researcher as applicable. Metabolite identity was established using a combination of an in-house metabolite library developed using pure purchased standards and the human metabolite data base (hmdb.ca). Isotope correction was performed using in-house software.

#### Mitochondrial phospholipids enrichment

Isolated mitochondria (500 μg) from 2-month-old mice were incubated in fusion buffer [220 mM mannitol, 70 mM sucrose, 2 mM Hepes, 10 mM KH2PO4, 5 mM MgCl2, 1 mM EGTA, 10 mM glutamate, 2 mM malate, 10 mM pyruvate, and 2.5 mM ADP (pH 6.5)] for 20 min at 30°C under constant stirring agitation in the presence of 15 nmol of small unilamellar vesicles (SUVs) (CL from Avanti Polar Lipids, 840012; PC from Avanti Polar Lipids, 850375C). After fusion, mitochondria were layered on a sucrose gradient (0.6 M) and centrifuged 10 min at 10,000g at 4°C to remove SUV. Pellet was then washed in mitochondrial buffer [250 mM sucrose, 3 mM EGTA, and 10 mM tris-HCl, (pH 7.4)].

#### Serum AST and ALT

Mice were sacrificed by CO_2_ inhalation and blood samples collected via cardiac puncture into heparin-coated tubes and centrifuged for collection of plasma within 1 hour of blood collection and frozen at -80°C until analysis. Plasma samples from mice were processed in a single batch for determination of serum alanine aminotransferase (ALT) and aspartate aminotransferase (AST) levels using a DC Element chemistry analyser (HESKA).

#### Quantification and statistical analyses

The level of significance was set at p < 0.05. Student’s T tests were used to determine the significance between experimental groups and two-way ANOVA analysis followed by a within-row pairwise comparison post hoc test where appropriate. For Student’s T tests, Gaussian distribution was assumed as well as assuming both populations had the same standard deviation. The sample size (n) for each experiment is shown in the figure legends and corresponds to the sample derived from the individual mice or for cell culture experiments on an individual batch of cells. Unless otherwise stated, statistical analyses were performed using GraphPad Prism software.

## Supporting information

Supplemental Table 1, Legends for Supplemental Figures

## Materials availability

Plasmids utilized by this study are available from Sigma Aldrich. Mouse lines generated by this study may be available at personal request from the lead contact. No new reagents were created or used by this study.

## Data and code availability

Individual data are presented in the figures. All data are available upon request.

## Author Contributions

MJB and KF contributed to conceptualization, experimental design, investigation, formal analysis, writing the original draft, and reviewing and editing the manuscript. GC, ADP, LW, and RHC performed investigation and formal analysis. JAM, QJP, and JMJ were involved in investigation, methodology, and formal analysis. AMP, TBB, JLS, SW, ERM, AMM, and STD contributed to investigation. PS, FMF, and ZGH contributed to experimental design, with PS and FMF also contributing to investigation, and ZGH contributing to resources and methodology. AP, SAP, GLH, and PNM were involved in investigation and methodology, with SAP also contributing to formal analysis. KEA and KJE contributed to investigation and resources. TST and MP contributed to conceptualization, with TST also contributing to investigation. LSN, SMN, DSL, and KHF-W contributed to methodology, with LSN also contributing to investigation. JLC contributed to methodology and formal analysis. WLH and SAS contributed to experimental design and resources. JEC contributed to methodology. All authors read and approved the paper.

Corresponding Author Correspondance to Katsuhiko Funai

## Ethics Declarations

Competing interests: the authors declare no competing interests.

---

**Figure 1– figure supplement 1.**
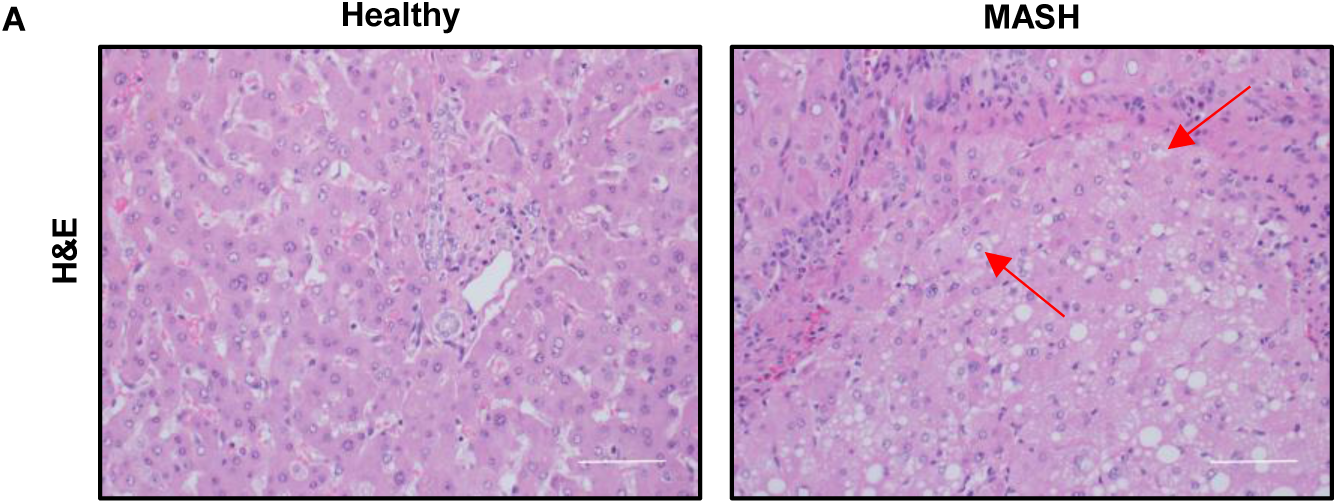

**Figure 1– figure supplement 2.**
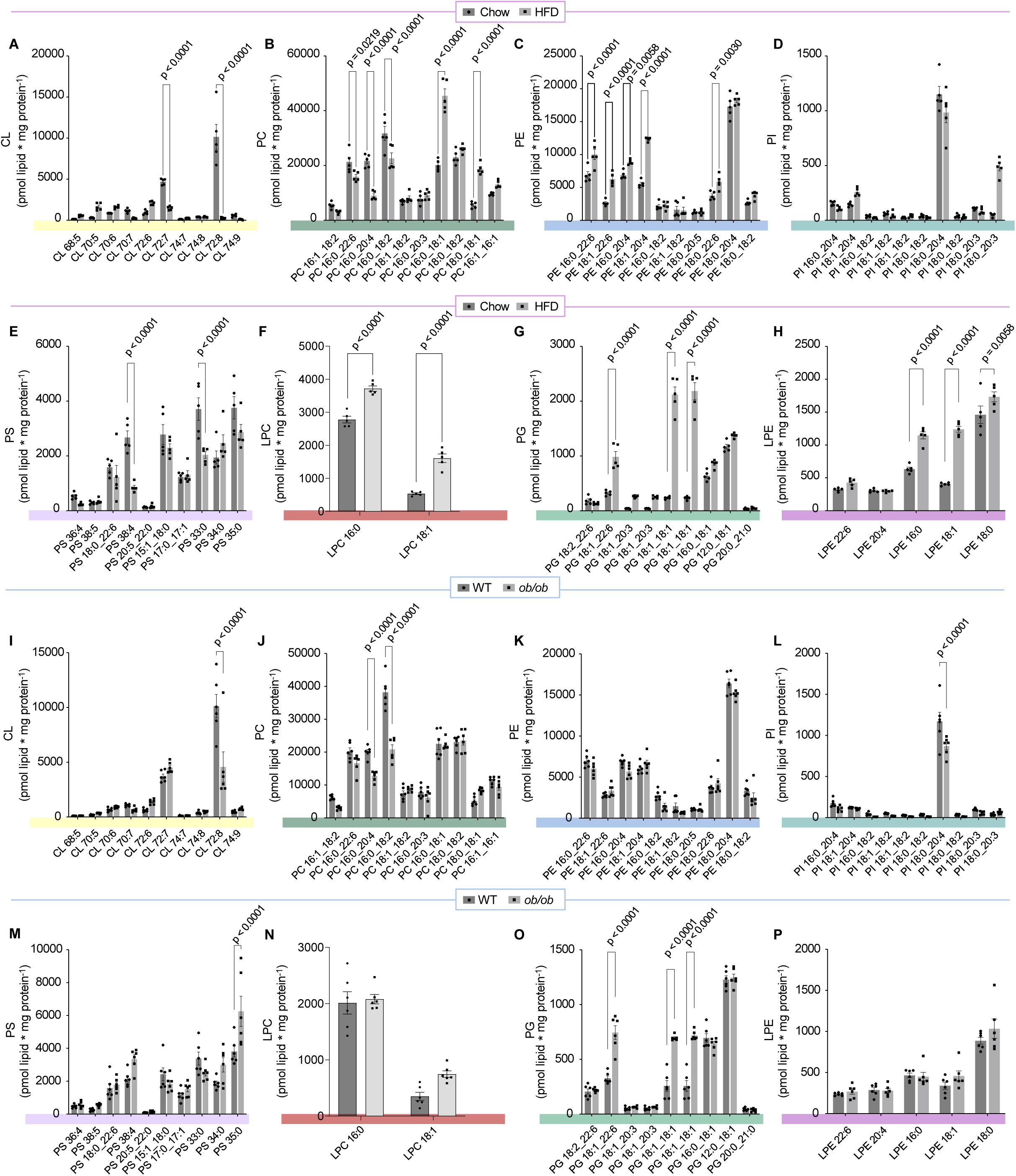

**Figure 1– figure supplement 3.**
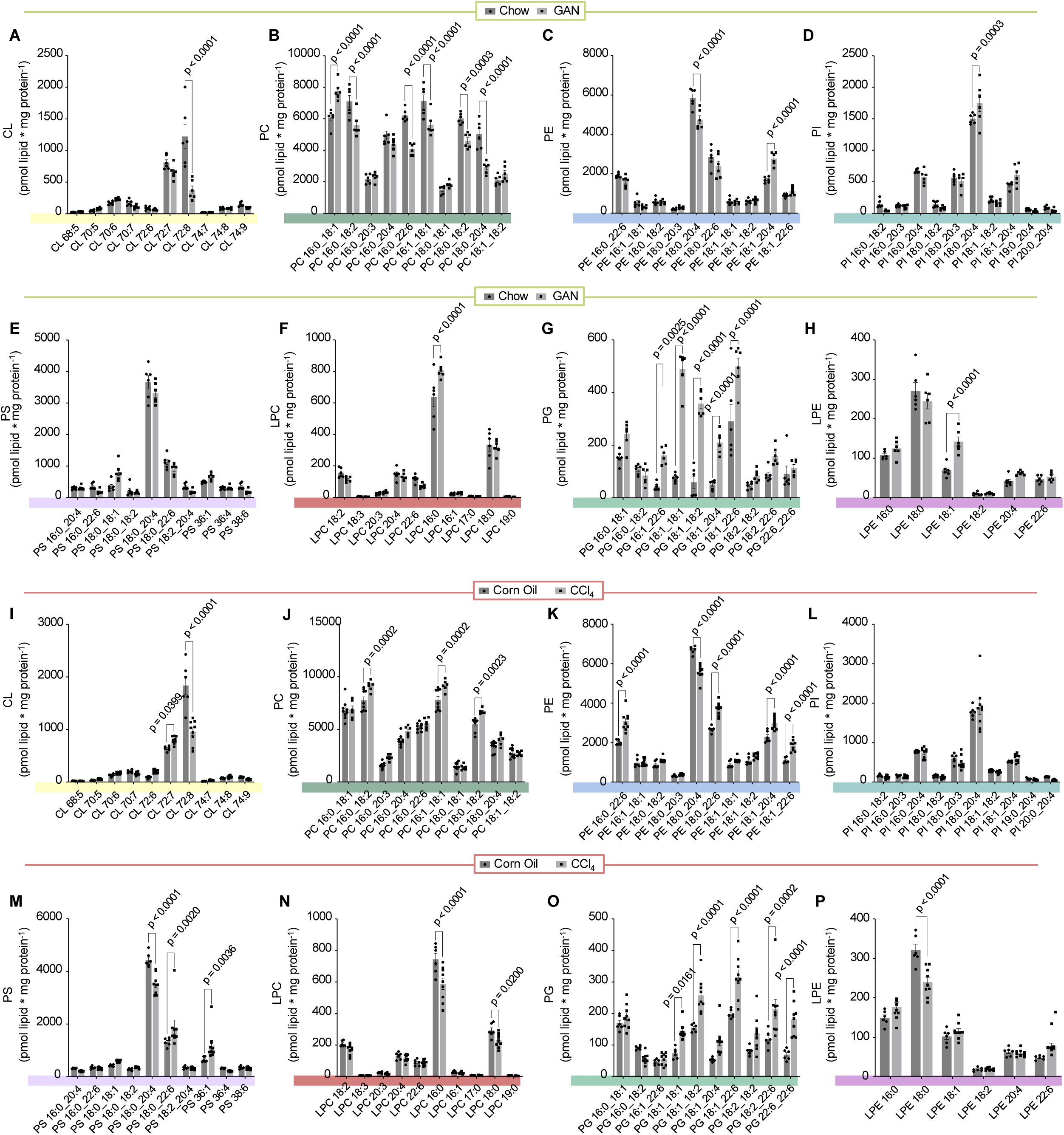

**Figure 2– figure supplement 1.**
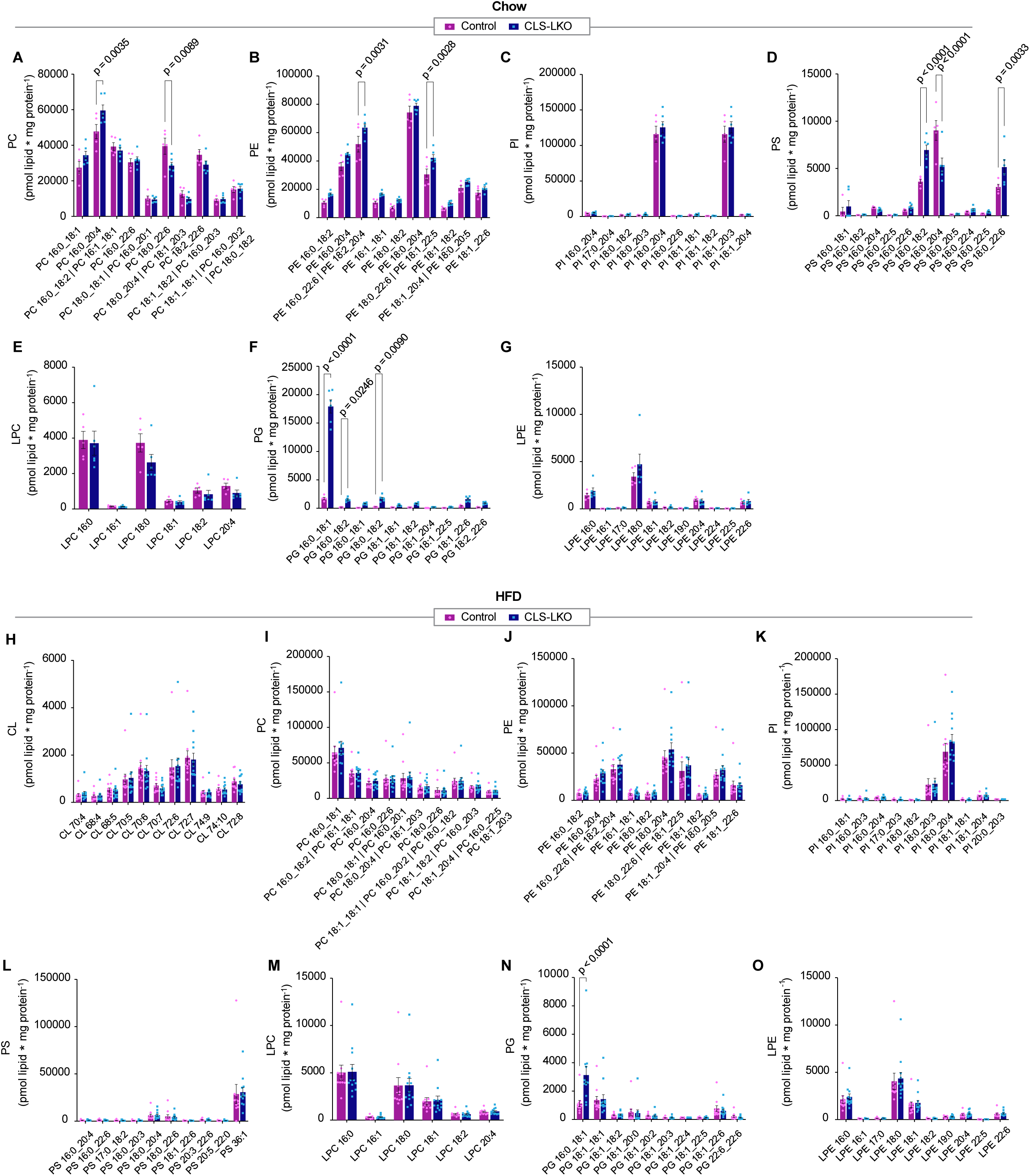

**Figure 2– figure supplement 2.**
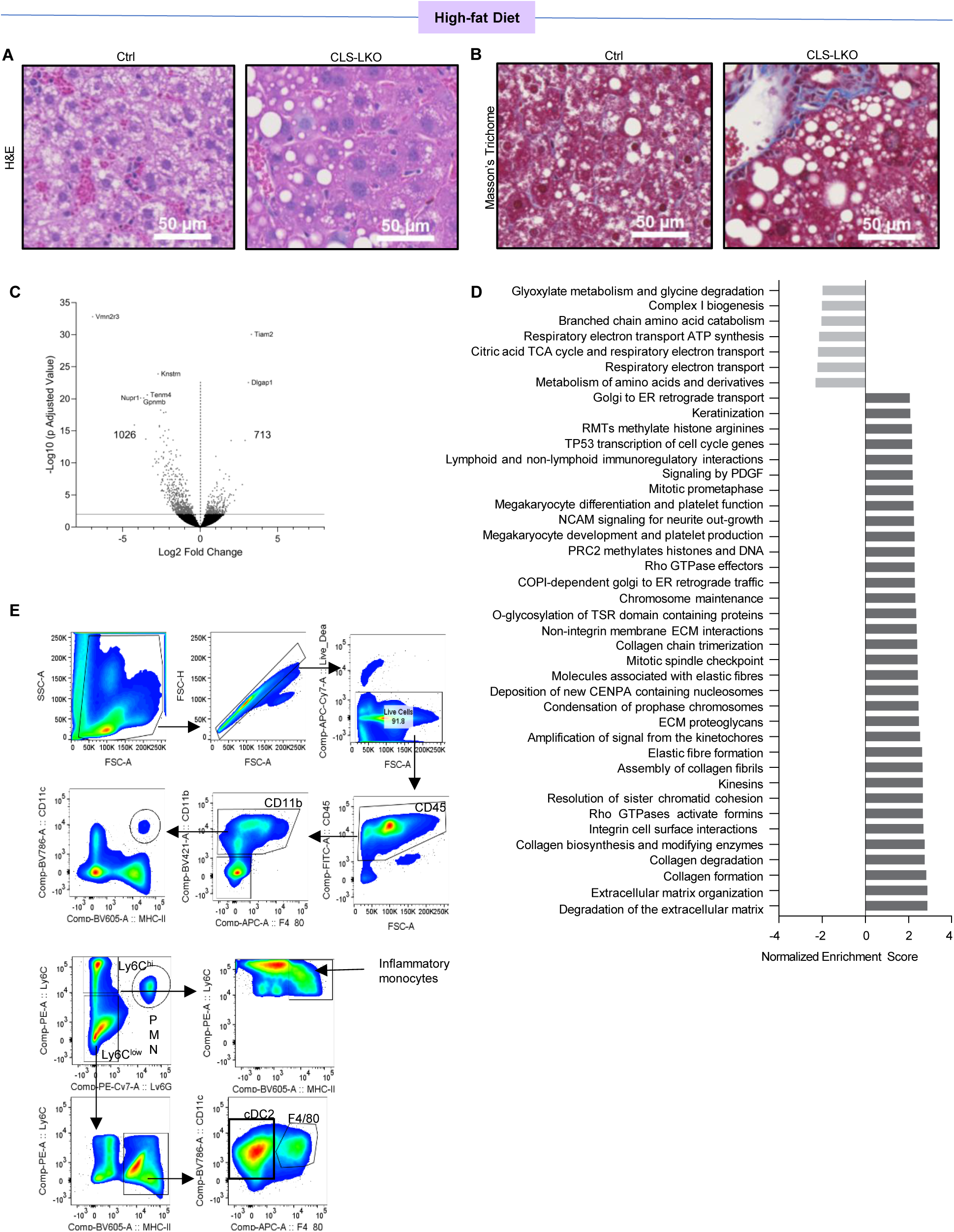

**Figure 3– figure supplement 1.**
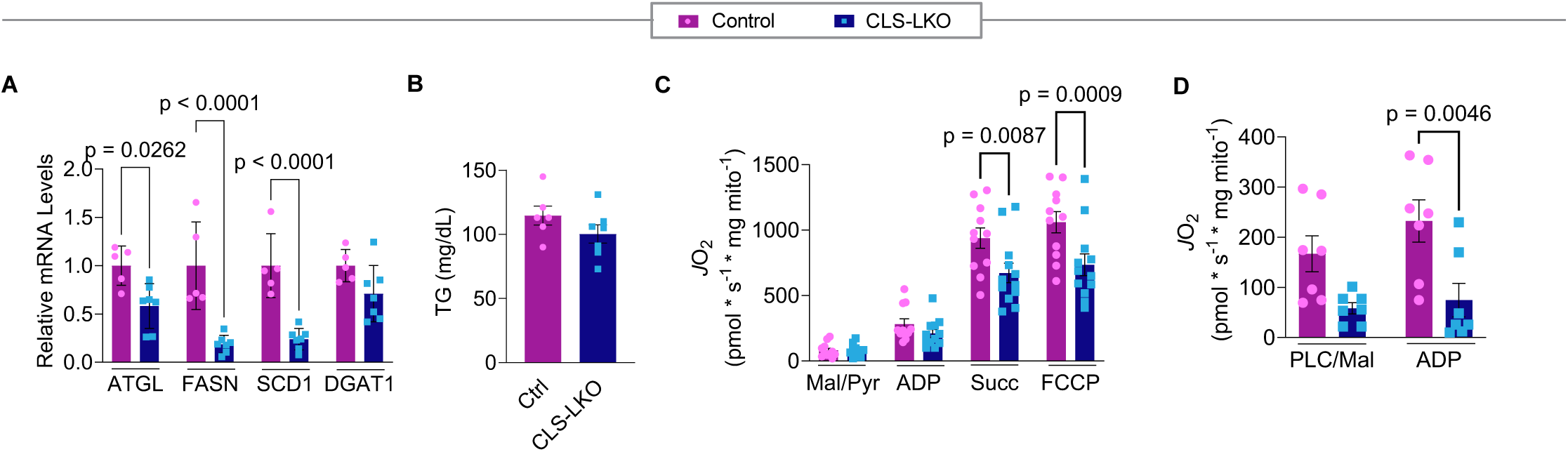

**Figure 4– figure supplement 1.**
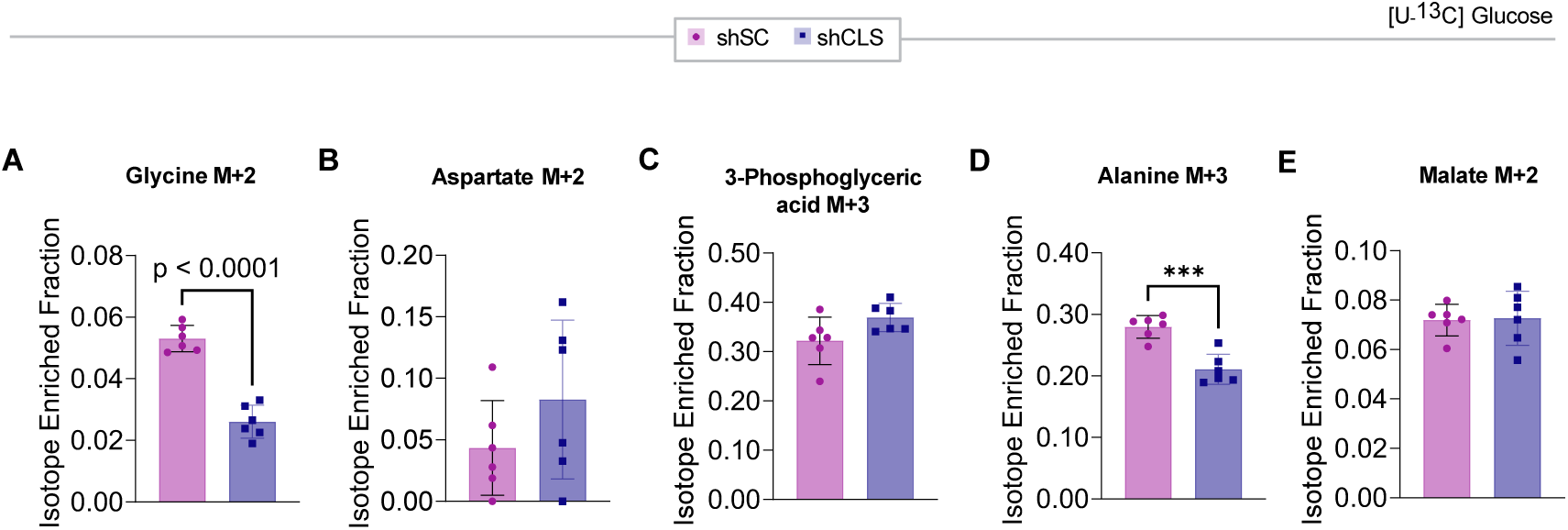

**Figure 5– figure supplement 1.**
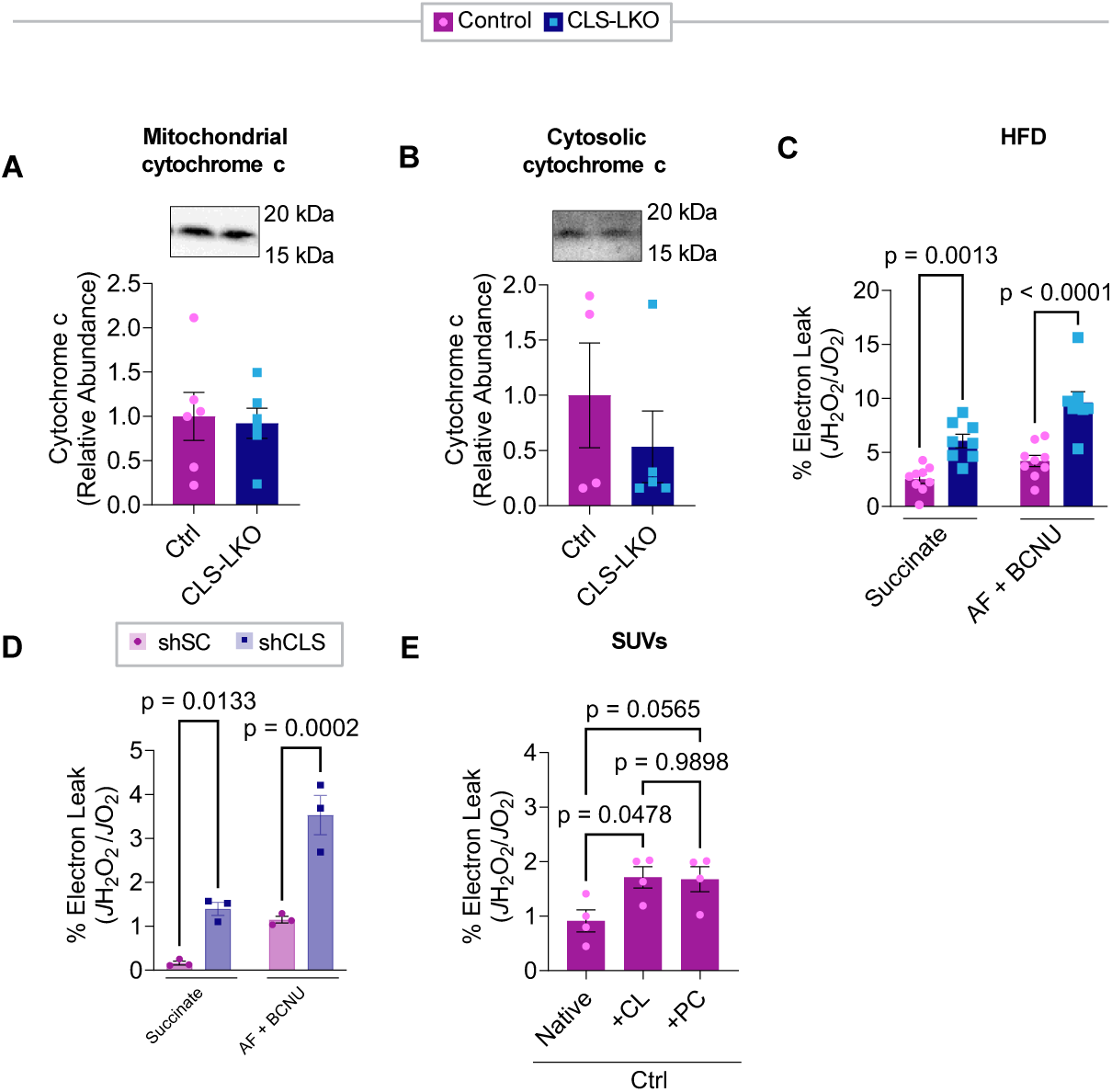

**Figure 7– figure supplement 1.**
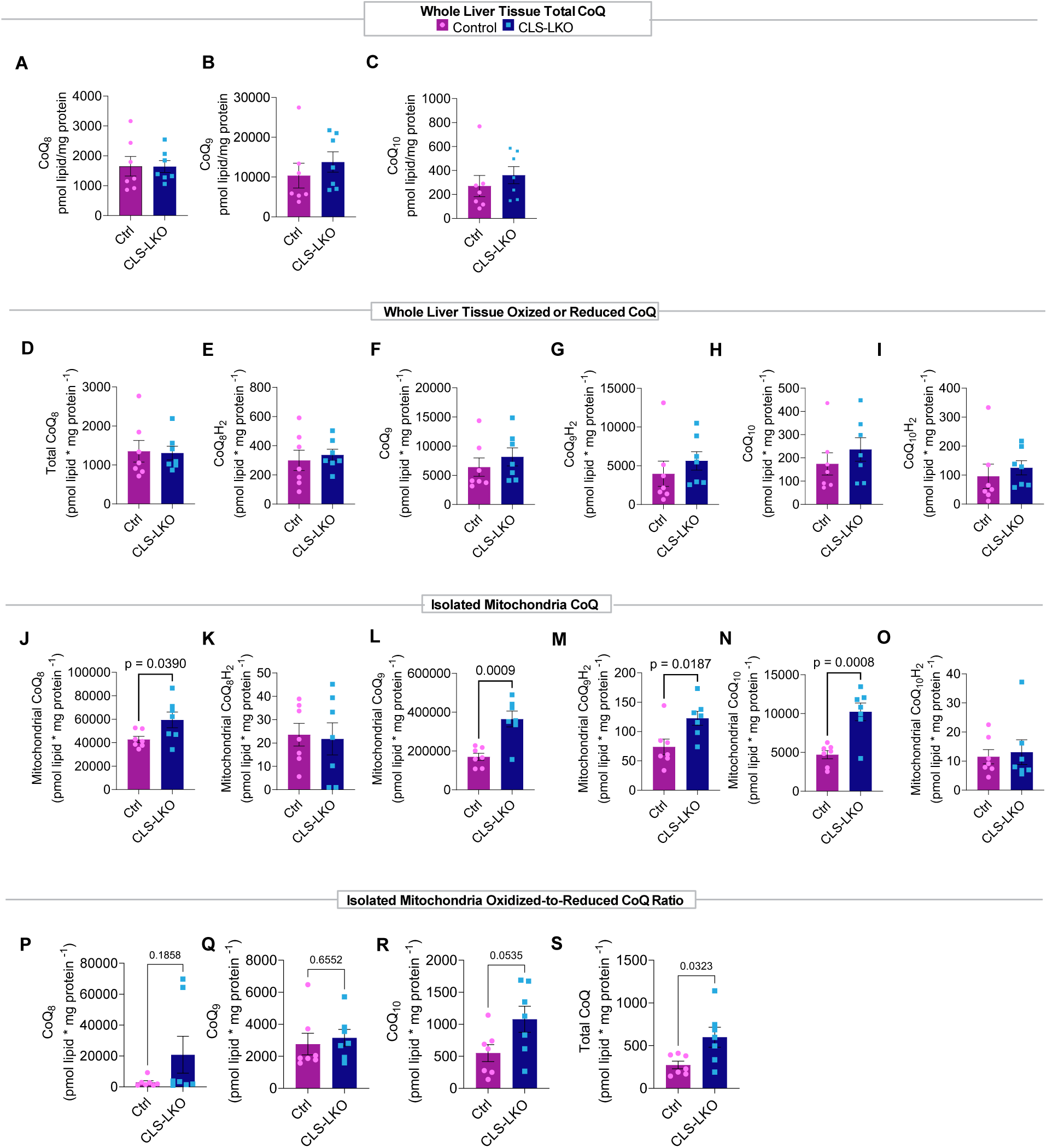

## Notes

### Competing Interest Statement

The authors have declared no competing interest.

### Summary of Updates

The manuscript was peer-reviewed at eLife. This revised manuscript submitted as revision to eLife includes new experiments, new data, modified text compared to the previous version.

### Funding

- National Institute of Diabetes and Digestive and Kidney Diseases: DK107397, DK127979
- National Institute on Aging: AG074535
- National Institute of General Medical Sciences: GM144613

