## Supplemental Table 1, Legends for Supplemental Figures for "Cardiolipin deficiency disrupts electron transport chain and drives steatohepatitis"

**Figure 1– table supplement 1**

Patient demographic information

|  | Healthy | MASH |
| --- | --- | --- |
| Age at time of collection | 50.3 ± 9.6 yrs | 62.2 ± 7.1 yrs |
| Sex | Male: 1 Female: 10 | Male: 9 Female: 8 |
| Alcohol use? | N/A | Yes: 0 No: 17 |
| Race | White: 9 African American: 1 Asian: 1 | White: 10 Unknown: 7 |

**Supplemental Figure Legends**

**Figure 1 – figure supplement 1.** (A) Representative H&E-stained liver sections from a healthy human subject (left) and a patient with MASH (right) at 20x magnification; red arrows indicated ballooned hepatocytes.

**Figure 1 – figure supplement 2. Mitochondrial phospholipidome from Figure 1N&O.** (A-H) Abundance of mitochondrial lipids (CL, PC, PE, PI, PS, LPC, PG, LPE) in livers of mice fed standard chow or HFD for 16 weeks (n=5 per group). (I-P) Abundance of mitochondrial lipids (CL, PC, PE, PI, PS, LPC, PG, LPE) in livers of wildtype or leptin-deficient mice, 30 weeks old (n=6 per group). Statistical significance was determined by 2-way ANOVA with within-row pairwise comparison. Data represent mean ± SEM. All measurements were taken from distinct samples.

**Figure 1 – figure supplement 3. Mitochondrial phospholipidome from Figure 1P&Q.** (A-H) Abundance of mitochondrial lipids (CL, PC, PE, PI, PS, LPC, PG, LPE) in livers of mice fed standard chow or Gubra-Amylin MASH diet for 30 weeks (*n*=6 per group). (I-P) Abundance of mitochondrial lipids (CL, PC, PE, PI, PS, LPC, PG, LPE) in livers of mice injected with corn oil or carbon tetrachloride for 10 weeks (*n*=5–7 per group). Statistical significance was determined by 2-way ANOVA with within-row pairwise comparison. Data represent mean ± SEM. All measurements were taken from distinct samples.

**Figure 2– figure supplement 1. Mitochondrial phospholipidome from standard chow or high-fat diet fed control and CLS-LKO livers.** (A-G) Abundance of mitochondrial lipids (PC, PE, PI, PS, LPC, PG, LPE) in livers of control or CLS-LKO mice fed standard chow, 8 weeks old (*n*=5–6 per group). (H-O) Abundance of mitochondrial lipids (CL, PC, PE, PI, PS, LPC, PG, LPE) in livers of control or CLS-LKO mice fed a high-fat diet for 8 weeks (*n*=11–12 per group). Statistical significance was determined by 2-way ANOVA with within-row pairwise comparison. Data represent mean ± SEM. All measurements were taken from distinct samples.

**Figure 2– figure supplement 2. Additional histological and transcriptomic data from control and CLS-LKO mice.** (A) Representative H&E staining of livers from control or CLS-LKO mice fed a high-fat diet (HFD) for 8 weeks. (B) Representative Masson’s Trichrome staining of livers from control or CLS-LKO mice fed a HFD for 8 weeks. (C) Volcano plot of differentially expressed genes in livers from control or CLS-LKO mice fed standard chow (*n*=5–7 per group). (D) Pathway analysis of transcriptomic data in livers from control or CLS-LKO mice fed standard chow (*n*=5–7 per group). (E) Gating strategy used for flow cytometry experiments in control and CLS-LKO mice. Pathway analysis was performed using the Reactome Pathway Database. All measurements were taken from distinct samples.

**Figure 3– figure supplement 1. Additional metabolic and mitochondrial phenotyping data with CLS deletion.** (A) mRNA levels of lipogenic genes in livers from control or CLS-LKO mice fed standard chow (*n*=6–7 per group). (B) Serum triglycerides in control or CLS-LKO mice fed standard chow (*n*=6–7 per group). (C,D) *J*O₂ consumption in isolated liver mitochondria from control or CLS-LKO mice fed a high-fat diet (HFD) for 8 weeks in response to substrates: (C) malate, pyruvate, ADP, succinate, and FCCP (*n*=11–12 per group); (D) palmitoyl-carnitine, malate, and ADP (*n*=7 per group). Statistical significance was determined by 2-way ANOVA with within-row pairwise comparison (A,C,D) and unpaired Student's t-test (B). Data represent mean ± SEM. All measurements were taken from distinct samples.

**Figure 4– figure supplement 1. Additional fluxomic phenotyping data with CLS deletion.** (A-D) Levels of labeled metabolites (glycine, aspartate, 3-phosphoglyceric acid, alanine, and malate) from glucose tracing in Hepa1-6 cells without (shSC) or with CLS deletion (shSLC) (*n*=6 per group). Statistical significance was determined by unpaired Student's t-test. Data represent mean ± SEM. All measurements were taken from distinct samples.

**Figure 5– figure supplement 1. Additional mitochondrial phenotyping data with CLS deletion.** (A,B) Levels of mitochondrial and cytosolic cytochrome c in livers of control or CLS-LKO mice fed a high-fat diet (HFD) for 8 weeks (*n*=6 per group). (C,D) *J*H₂O₂ emission and production in isolated liver mitochondria from (C) control or CLS-LKO mice fed a HFD for 8 weeks, stimulated with succinate or succinate plus auranofin and BCNU (*n*=9–8 per group), and (D) Hepa1-6 cells without (shSC) or with CLS deletion (shCLS), stimulated with succinate or succinate plus auranofin and BCNU (*n*=3 per group). (E) Electron leak of liver mitochondria from control mice fused with SUVs (*n*=4 per group). Statistical significance was determined by 2-way ANOVA with within-row pairwise comparison (C,D,E) and unpaired Student's t-test (A,B). All measurements were taken from distinct samples.

**Figure 7– figure supplement 1. Additional data on coenzyme Q** (A-C) Total CoQ_8_, CoQ_9_, and CoQ_10_ levels in whole liver tissue from control and CLS-LKO mice (*n*=7 per group). (D-I) Oxidized and reduced CoQ_8_, CoQ_9_, and CoQ_10_ levels in whole liver tissue from control and CLS-LKO mice (*n*=7 per group). (J-O) Oxidized and reduced CoQ_8_, CoQ_9_, and CoQ_10_ levels in isolated mitochondria from control and CLS-LKO mice (*n*=7 per group). (P-S) Oxidized-to-reduced CoQ ratios in isolated liver mitochondria for CoQ_8_, CoQ_9_, CoQ_10_, and total CoQ calculated from the same samples as in panels J-O. Statistical significance was determined by unpaired Student's t-test. Data represent mean ± SEM. All measurements were taken from distinct samples.
